# Virus-like antigen display delivers a stand–alone danger signal through the BCR that circumvents tolerance

**DOI:** 10.64898/2026.04.23.720482

**Authors:** Julianne B. Riggs, Alexander J. Ritter, François X. P. Bourassa, Alexander R. Meyer, Erika Kay-Tsumagari, Wei-Yun Wholey, Jeremy Libang, James L. Mueller, Wen Lu, Ned S. Wingreen, Wei Cheng, Julie Zikherman

## Abstract

How B cells discriminate self from foreign antigens remains a central question, given inherent autoreactivity of the mature B cell receptor (BCR) repertoire. Soluble antigen (sAg) induces tolerance, whereas patterned antigen display on virus-like particles (pAg) triggers robust B cell responses that can proceed without T cell help. Here, we show how this divergence arises early in BCR signaling. Unlike sAg, pAg can bypass a Lyn-dependent negative feedback loop to trigger digital signaling, such that ultra-low concentrations of pAg produce strong and sustained Ca^2+^ responses. Surprisingly, pAg drives maximal nuclear NF-κB but limited NFAT, whereas sAg does the opposite, reflecting differential production of diacylglycerol. Consequently, sAg induced an NFAT-dependent anergy program, whereas pAg evaded this state and instead engaged a cMyc-driven program that partially resembles a TLR-dependent danger response. Our findings reveal how proximal signaling directs distinct transcriptional fate to enable immunogenic B cell responses to virus-like antigen display.

## Introduction

Some pathogens, including certain viruses, are shaped by evolution to acquire self-like features that enable host-cell infection while evading immune recognition ^1,2^. Self-reactivity is a normal feature of the mature B cell repertoire and may protect the host against such self-mimicking pathogens ^3–8^. In support of this model, protective antibodies against vaccinia, SARS-CoV-2, and HIV can originate from germ-line-encoded, self-reactive B cell antigen receptors (BCRs) ^9–15^. However, retaining rather than eliminating self-reactive BCRs confers risk of autoimmunity. Indeed, molecular mimicry by viruses can trigger humoral autoimmunity^1^. Yet, most healthy individuals are able to both clear these pathogens and evade pathogenic self-targeting immune responses, successfully discriminating self from foreign. How B cells are wired to achieve this is not fully understood.

BCR signal transduction is tightly controlled by a balance of activating and inhibitory co-receptors that converge at the level of the second messenger PI(3,4,5)P_3_ (PIP_3_ herein) which nucleates assembly of a “signalosome” at the plasma membrane ^16,17^. Dynamic regulation of PIP_3_ links proximal BCR signaling to downstream pathways, including store-operated Ca^2+^ entry and diacylglycerol (DAG)-dependent activation of MAPK cascades and NF-κB. The activating co-receptor CD19 amplifies PIP_3_ production by recruiting PI3K, while a negative feedback loop, operated by the Src family kinase Lyn, engages inhibitory co-receptors such as CD22 that in turn recruit phosphatases to degrade PIP_3_ ^18^. The balance of these two arms influences whether a B cell will be unleashed for robust humoral immunity or restrained. Defects in the engagement or expression of Lyn, CD22, other inhibitory co-receptors, and their effector phosphatases are linked to autoimmune phenotypes in mice and disease in humans ^18–22^.

Soluble antigens (sAg) have long been used to study B cell tolerance to self ^23^. Without recruitment of additional co-stimulation in the form of T cell help or a pathogen-associated danger signal, sAg stimulation of the BCR produces, at most, an abortive B cell response – “signal 1 without signal 2” ^24^. If such help is not recruited within a limited time window, the outcome is tolerance characterized by functional unresponsiveness (“anergy”) and/or eventual apoptosis ^23,25^. B cell anergy is maintained by inhibitory co-receptors and phosphatases that constrain PIP_3_ abundance ^22,26,27^, and further enforced by downstream transcription factors ^28,29^. Imbalanced nuclear translocation of the Ca^2+^-dependent transcription factor NFAT in the absence of AP-1 has been well-recognized for its role in T cell anergy ^30^, and this pathway has been implicated in B cell anergy as well ^31–35^.

By contrast, it has long been appreciated that B cell responses to highly multivalent antigens, such as viruses or virus-like structures, are extremely rapid and robust, and do not require T cell help ^36–39^. These characteristics are critical for mounting immune responses when time is of the essence in the context of viral infection. Virus-like antigenic display has been harnessed to generate successful vaccines (e.g. HPV, Hepatitis B). Naturally-occurring viruses and virus-like formats vary greatly in their exterior and interior composition, but regular spacing and patterned display of antigenic epitopes is an essential feature ^40^. Moreover, this feature is encountered and recognized directly by B cells through their BCRs as intact particulate Ag (pAg) and viruses can traffic via lymph or blood to secondary lymphoid organs ^41–46^. However, the mechanism by which this fundamental biophysical feature of virus-like antigen display produces rapid, robust, and T-independent B cell responses is not fully understood.

Using synthetic virus-like structures (SVLS), a new generation of well-characterized pAg that have been validated both in vitro and in vivo for immunogenicity ^47,48^, we previously demonstrated that virus-like Ag display is a vastly more potent B cell stimulus than the same Ag in soluble form ^49^. Tethering Ag to a liposome of viral size produced robust BCR signaling responses that proceed in the absence of co-stimulation, due in part to evasion of Lyn-dependent inhibitory signals. By contrast, sAg was constrained by inhibitory tone and dependent upon T cell help. However, the molecular mechanisms linking distinct early BCR signaling by virus-like and soluble Ag with divergent downstream fates remain to be defined.

Here we continue to leverage viral-sized liposomal particles decorated with a model antigen to dissect how naïve murine B cells respond to biophysical formats of Ag. We show that pAg but not sAg trigger digital signaling responses by selectively evading a Lyn-dependent negative feedback loop. This is partially accounted for by the Lyn substrate CD22. Live cell imaging with fluorescently-labeled particles reveals heightened and prolonged calcium responses at the single cell level, and kinetics consistent with B cell activation by a very low number of particles. Of note, we find that pAg does not merely produce stronger downstream B cell activation than sAg, but rather qualitatively distinct and biased patterns of NFAT and NF-κB nuclear translocation, attributable to high DAG concentration. Consequently, sAg produces an NFAT-dependent anergy program while pAg largely evades this and instead induces a c-Myc-driven program to support cell growth and clonal expansion, even in the absence of T cell help. Nevertheless, pAg-stimulated B cells are able to efficiently activate T cells in an Ag-specific manner. Importantly, we reveal that pAg does not simply mimic the effects of CD40 co-stimulation, but rather delivers a unique signal through the BCR that recapitulates some transcriptional features of classic innate pattern recognition receptors. Our work reveals how virus-like antigen display delivers a stand-alone “danger signal” to B cells through the BCR to evade tolerance normally triggered by signal 1 in the absence of signal 2.

## Results

### Virus-like antigen produces digital BCR signaling by evading a Lyn-dependent negative feedback loop

To study B cell responses to virus-like antigen display, we took advantage of virus-sized liposomes (120nm diameter) composed of physiological lipids and decorated with model antigen at programmable density via maleimide-thiol coupling (**Figure 1A**) ^48,49^. This modular platform uses highly purified, well-defined ingredients assembled in vitro. We selected hen egg lysozyme (HEL) as a model antigen to pair with HEL-specific Hy10 BCR Tg (“MD4”) mice ^50^. Because the Hy10 BCR has extremely high, nanomolar affinity for native HEL protein (K_a_ ∼10^10^ M^-1^), we generated liposomes conjugated to HEL protein variants harboring affinity-reducing point mutations in the binding interface: HELD (R73E, D101R; K_a_ ∼10^8^ M^-1^) and HELT (R21Q, R73E, D101R; K_a_ ∼10^6^-10^7^ M^-1^) **(Figure S1A)**^51–54^.

**Figure 1:**
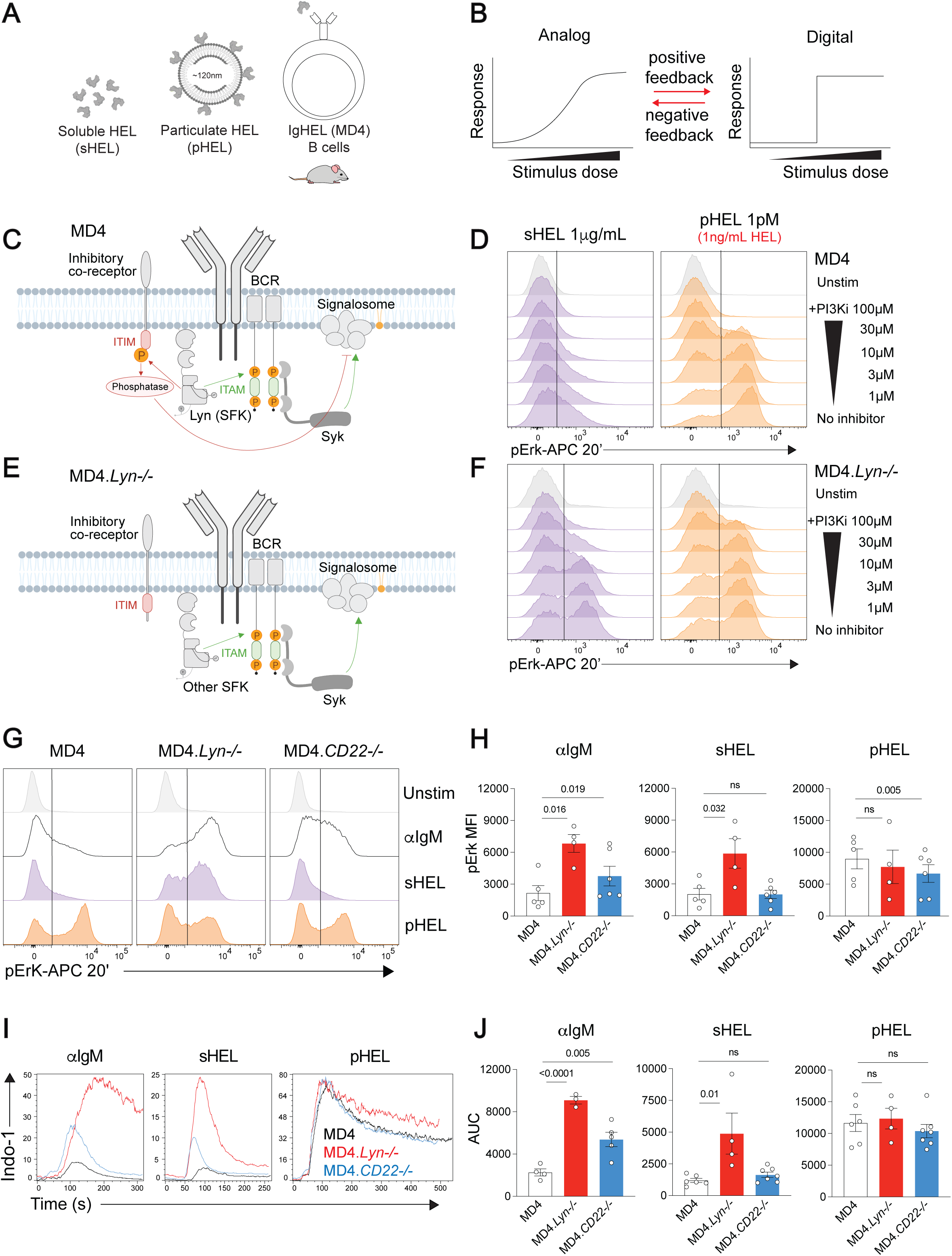
Virus-like antigen produces ultrasensitive, digital BCR signaling by evading a Lyn-dependent negative feedback loop 1. **A:** Toolkit for studying B cell responses to soluble and viral-like antigen. Particles composed of neutral lipids and HEL protein conjugated at high density (pHEL) were compared to soluble HEL (sHEL) by stimulating HEL-specific B cells from the MD4 mouse line that harbor the Hy10 BCR Tg. **1B:** Schematic depicts analog and digital signal responses; analog responses scale with stimulus dose, while digital responses exhibit a discrete switch between no response and maximal response at a particular stimulus dose. Digital signaling can be achieved by engagement of positive feedback and/or evasion of negative feedback. **1C:** Model of proximal BCR signal transduction in WT, Lyn-sufficient B cells. Lyn non-redundantly mediates ITIM phosphorylation and recruitment of inhibitory phosphatases to suppress downstream signaling, engaging negative feedback control. **1D:** Intracellular phospho-Erk (pErk) induction measured by flow cytometry 20 minutes post stimulation. MD4 pooled splenocytes and lymph node cells were pre-incubated with a dose titration of PI3K inhibitor Ly294002 (100, 30, 10, 3 and 1 μM) followed by stimulation with sHEL (1μg/mL) or pHEL (1pM). Histograms depict pErk gMFI of live B220+ cells and are representative of at least three independent experiments. **1E:** Model of signaling in Lyn-deficient B cells, featuring loss of coupling to ITIM inhibitory receptors and consequent elimination of negative feedback. **1F:** pErk measurement as in **1D** but using MD4.*Lyn−/−* splenocytes. **1G:** MD4, MD4.*Lyn−/−* and MD4.*CD22−/−* pooled splenocytes and lymph node cells were stimulated at 37C for 20 minutes and assessed for intracellular pErk by flow cytometry. Stimulus doses: 10μg/mL anti-IgM, 1μg/mL sHEL or 1pM pHEL. Histograms depict pErk gMFI in B220+ cells and are representative of six independent experiments. **1H:** Quantification of data in **1G**. Each data point represents pErk-APC mean fluorescence intensity (MFI) of B220+ cells from one experiment. P-values acquired via one-way ANOVA. **1I:** MD4, MD4.*Lyn−/−* and MD4.*CD22−/−* pooled splenocytes and lymph node cells were loaded with Indo-1 Ca^2+^ indicator dye and stained for viability and surface markers. Cells were run in real time by flow cytometry to assess intracellular Ca^2+^ entry after stimulation with ionomycin 1μM, anti-IgM 10μg/mL, sHEL 1μg/mL or pHEL 1pM. Plots depict geometric mean of Indo-1 bound/unbound fluorescence ratio in individual live B220^+^ CD23^+^ cells over time (seconds). Data representative of three independent experiments. **1J:** Quantification of data in **1I**. Area under the Ca^2+^ mobilization curves. Each data point represents an independent experimental replicate. P-values acquired via one-way ANOVA.

We previously used this library of antigen-conjugated liposomes to demonstrate that naïve B cell responses to pAg display are highly potent, density-dependent, and affinity-independent ^49^. For the purposes of the present study, we selected liposomes with a high epitope density of HEL (>100 proteins/particle). Additionally, we use particles with HELD and HELT (particulate HEL or pHEL) interchangeably, as we previously found that B cell responses to these variants was indistinguishable ^49^. As a model sAg comparator – except where explicitly noted – we used native HEL in soluble form (sHEL) rather than low affinity HEL variants in order to trigger B cell signaling robust enough to be assayed. For all experiments we selected equipotent doses of pHEL and sHEL established by prior studies ^49^ which generate comparable activation of the BCR signaling reporter Nur77-eGFP, but correspond to almost 1000x difference in HEL protein concentration: sHEL (1μg/mL) and pHEL (1pM, ∼1ng/mL HEL protein).

We compared proximal BCR signaling responses of Ag-specific MD4 B cells to sHEL and pHEL. We previously observed that even very high doses of sHEL induced modest Erk phosphorylation (pErk), while near-maximal pErk was triggered by extremely low concentrations of pHEL ^49^. Importantly, both were suppressed by inhibitors of canonical nodes of the BCR signaling pathway: Syk, Btk, and PI3K (**Figures S1B, C, D**) ^49^. Syk can phosphorylate immunotyrosine activating motifs (ITAMs) in the absence of Src family kinases (SFKs) in response to BCR cross-linking by a multivalent Ag ^55^. However, B cells show no response to pHEL when treated with a pan-SFK inhibitor (PP2) (**Figure S1B, E**).

All-or-none, bimodal responses at the population level imply digital signal transduction (**Fig 1B**)^56,57^. Erk phosphorylation assayed in single cells responding to pHEL – but not sHEL – exhibits a bimodal distribution, with two distinct peaks corresponding to non-responding cells and maximal responders (**Figure 1C, D**). To test for true bimodality of this response to pHEL, we broadly titrated concentration of the PI3K inhibitor Ly294002. As noted, high doses of inhibitor completely block Erk phosphorylation in response to pHEL. As inhibitor concentration is decreased, the fraction of responders increases, but bimodality persists (**Figure 1D**). This provides rigorous evidence for bona fide digital signaling by pHEL stimulation. In contrast, the population distribution of pErk response to sHEL under PI3K inhibition remains unimodal/continuous, implying analog rather than digital signal transduction. These patterns were preserved with titration of both Syk and Btk inhibitors (**Figure S1B-D**).

Digital signal transduction is typically achieved by engagement of a positive feedback loop and/or evasion of negative feedback (**Figure 1B**)^56,57^. B cells are wired to have both, balancing both activating and inhibitory arms (**Figure 1C**). At the nexus of these arms is the SFK Lyn ^18^. Lyn plays a redundant role with other SFKs in phosphorylation of BCR ITAMs but a *non-redundant* function in phosphorylation of immunotyrosine inhibitory motifs (ITIMs) on inhibitory co-receptors. ITIM phosphorylation facilitates dynamic recruitment and activation of protein tyrosine and lipid phosphatases that suppress the BCR signal transduction cascade ^19^. Deletion of Lyn therefore selectively ablates ITIM-dependent signaling but leaves ITAM-signaling intact (**Figure 1E**).

We previously showed that sHEL but not pHEL responses were constrained by Lyn and exhibit signal amplification at the level of PIP_3_ ^49^. This suggested that potent B cell responses to pHEL were due in part to evasion of Lyn-dependent inhibitory signaling. Indeed, in the absence of Lyn, MD4 B cells exhibit increased pErk in response to sHEL, but no further boost in response to pHEL **(Figure 1F)**. Intriguingly, we now show that in Lyn-deficient B cells, pErk responses to sHEL exhibit bimodal population distribution and dose-titration of PI3K inhibitor preserves this bimodality **(Figure 1F)**. This suggests that evasion of Lyn-dependent inhibitory tone provides not only a mechanistic explanation for the potency gap between sHEL and pHEL, but also accounts for digital signaling responses to pHEL. Interestingly, we noted that B cell responses to pHEL exhibit unique sensitivity to Btk inhibition (**Fig S1D**). This contrasts with more proximal pathway inhibitors (Syk and PI3K). Btk represents a potential node of signal amplification in pHEL-activated B cells following recruitment via PIP_3_ to the membrane and dimerization, which may account for this observation ^58^.

### Negative feedback by Lyn is partially mediated by CD22

We next sought to identify Lyn substrates that may mediate suppression of B cell responses to soluble BCR stimuli but not virus-like Ag display. CD22 is an ITIM-containing inhibitory co-receptor that is closely associated with the IgM BCR and is excluded from the immunological synapse following encounter with Ag presented on membranes ^20,21,59^. CD22 facilitates tolerance by binding self-sialic acid ligands on host molecules, both in trans on neighboring cells and in cis on other transmembrane proteins, such as the IgM BCR ^20,21^. CD22 is therefore an important safeguard against autoimmunity, in both humans and mice ^21,60–63^.

To study the role of CD22 in B cell responses to soluble and particulate Ag, we generated MD4*.CD22−/−* mice and compared them to MD4 and MD4.*Lyn−/−* mice. These mice had relatively normal B cell numbers, by contrast to the significant B cell depletion seen in MD4.*Lyn−/−* (**Figure S1F**). pAg responses by MD4 B cells are not altered in the absence of either Lyn or CD22, consistent with evasion of the Lyn-CD22-inhibitory axis (**Figure 1G-J**). By contrast, deletion of CD22 enhances pErk and Ca^2+^ responses to anti-IgM Fab’2 stimulation (**Figure 1G-J**), albeit not to the extent observed in the absence of Lyn. However, sAg responses, while constrained by Lyn, are not significantly enhanced in the absence of CD22. We noticed profound downregulation of surface IgM on *CD22−/−* MD4 B cells, specifically in the CD23^hi^ CD21^int^ compartment (**Figure S1G-H**). We hypothesized that IgM downregulation by CD22−/− B cells could serve as a compensatory tolerance mechanism and may mask sHEL proximal signaling responses. To test this, we used IgM Fab’1 to mark B cells with distinct IgM surface expression without cross-linking the BCR. This indeed revealed that CD23+ IgM^hi^ MD4 B cells from CD22−/− mice exhibit heightened Ca^2+^ mobilization and Erk phosphorylation in response to sHEL (**Figure S1I-L**). We conclude that CD22 partially mediates Lyn-dependent inhibition of B cell responses to soluble Ag, but additional Lyn substrates must contribute.

### Particulate antigen induces sustained calcium responses at the single cell level

Flow cytometric analyses demonstrate heightened and prolonged cytosolic Ca^2+^ responses to pAg, as previously shown (**Figure 1I**)^49^. However, these data represent a population of cells sampled at serial time points and obscure single cell Ca^2+^ dynamics over time. Here we imaged Ca^2+^ responses with microscopy in live purified B cells loaded with Fluo4-AM Ca^2+^ indicator dye and affixed to a glass plate. To visualize particle-B cell interactions, we used a pHELD particle with AlexaFluor-594 as cargo (pHEL-AF594)^49^. We recorded images every two seconds for a total of five minutes following stimulation with either pHEL-AF594 or soluble BCR stimuli (**Figure 2A-D, Figure S2A, Videos S1-17**). From these recordings, we were able to generate single cell Ca^2+^ traces by tracking the Fluo4-AM intensity of individual B cells over time (**Figure S2B-C)**. We present representative videos and single cell traces of each stimulation condition as well as quantification across multiple imaged B cells.

**Figure 2:**
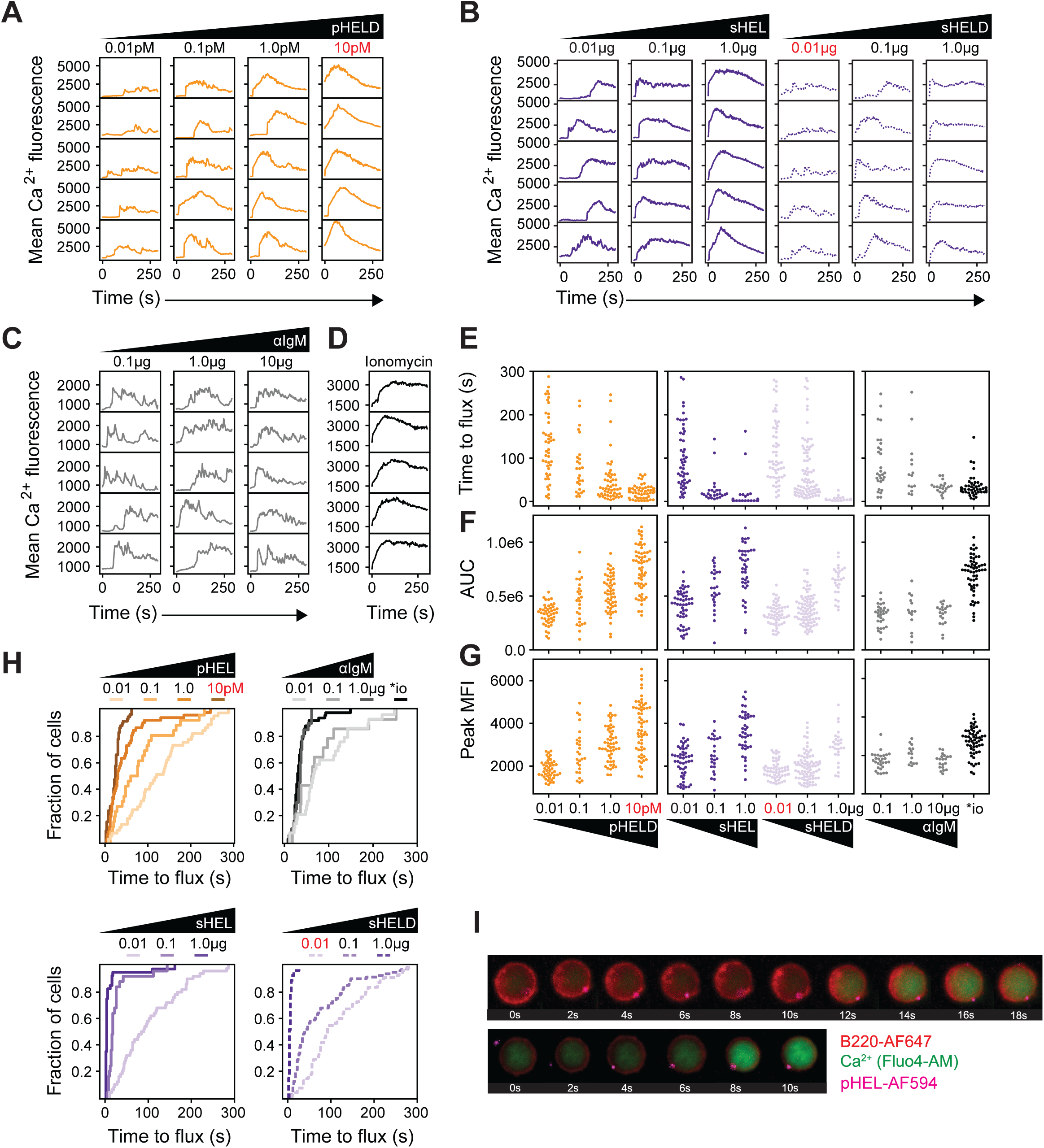
Particulate antigen induces sustained calcium flux at the single cell level 2A-D: Single cell traces of live B cells fluxing Ca^2+^ in response to dose titrations of pHELD-AF594 (**A**), sHEL, sHELD **(B**), anti-IgM **(C**), or ionomycin **(D)**, recorded via microscopy. Mean Ca^2+^ fluorescence intensity is plotted as a function of time. Traces were extracted from cells from supplementary videos and are representative of multiple B cells across three independent experiments. Red text denotes pHELD and sHELD doses with matched concentrations of HEL protein (sHELD 0.01ng/mL and pHELD 10pM). **2E:** Time to flux calculated from individual B cell traces. Flux was defined as a cell reaching 1.5x starting Ca^2+^ mean fluorescence intensity. **2F:** Area under the curve for individual B cell traces. **2G:** Peak Ca^2+^ fluorescence intensity for individual B cells. **2H:** Empirical cumulative distribution function (eCDF) showing the fraction of cells fluxing Ca^2+^ as a function of time across varying doses of stimulations. **2I:** Screenshots from Ca^2+^ videos of B cells stimulated with 10pM pHEL-AF594 showing encounters of single B cells with fluorescent particles.

We titrated pHEL-AF594 concentration across four orders of magnitude. A high dose of pAg (10pM) resulted in near synchronous increase of cytosolic Ca^2+^ with a high intensity peak fluorescence that resembled ionomycin response **(Figure 2A, 2D** and **Videos S1, S2**). Strikingly, we observed highly prolonged calcium responses (over many minutes) by individual B cells at 10pM and 1pM, suggesting that the long duration Ca^2+^ responses to pHEL detected by flow cytometry reflected the behavior of individual B cells rather than an asynchronous population average. As pAg concentration was reduced (1pM and 0.1pM), we observed asynchronous onset of Ca^2+^ response among imaged B cells with delayed activation kinetics in the population, as well as a progressive decline in integrated and peak Ca^2+^ concentration (**Figure 2A, E-H** and **Videos S3, S4**). At an ultralow particle concentration (0.01pM), most B cells displayed a detectable, but much lower intensity, Ca^2+^ response exhibiting oscillatory dynamics (**Figure 2A** and **Video S5**).

We next sought to compare these B cell responses to those triggered by sAg. Here we took advantage of HEL affinity variants (sHEL, sHELD) as well as broad dose titration. B cell responses to sAg showed marked sensitivity to both affinity and dose (**Figure 2B**, **2E-H** and **Videos S6-S11)**. We observed oscillatory behavior as dose was decreased, most evident with low affinity sHELD (**Figure 2B**). Importantly, comparing sAg and pAg highlights the large potency gap between these two types of stimuli; pHELD fluorescent particles at 1pM concentration induced a comparable peak and AUC Ca^2+^ as 1μg/mL sHELD, although the time to peak is longer. This reflects more than 500-fold lower concentration of HELD protein in particulate form, consistent with prior observations made using flow-based readouts ^49^.

Finally, we analyzed B cell responses to IgM cross-linking and observed striking calcium oscillations even at a high, saturating dose of anti-IgM Fab’2 (10μg/mL) (**Figure 2C** and **Videos S12-S14**). While time to activation scaled with anti-IgM dose titration, area under the curve and peak amplitude did not (**Figure 2E-G)**.

### Visualization and modeling of single particle B cell activation

Digital signaling responses to pAg imply that there is a threshold number of particles required for B cell activation (**Figure 1B, D, S1C**). Since extremely low concentrations of pHEL are sufficient to trigger B cell signaling, we sought to determine how many particles are required to activate an individual B cell. At the lowest concentration tested (0.01 pM), the particle-to-cell ratio was 1:3, suggesting that one viral particle may be sufficient to activate an individual B cell. Because our imaging studies enable us to simultaneously visualize fluorescent pHEL-AF594 particles and B cell calcium responses, we reasoned that this might present an opportunity to address this question. Indeed, we could identify in our calcium imaging videos examples of individual particle-B cell interactions followed by Ca^2+^ entry (**Figure 2I, Videos S15-S17**).

To gain mechanistic insight into the kinetics of pHEL-induced B cell activation, we analyzed cumulative distribution (CDF) plots depicting the fraction of activated B cells over time at varying pHEL concentrations (**Figure 2H**). We first fit the pHEL CDF plots with single exponential kinetics via maximum likelihood to examine how the observed rate constants (*k*_obs_) varied with pHEL concentration (**Figure S2D-E**). At low particle concentrations, *k*_obs_ values in our data set were close to the diffusion-limited binding rate estimated from first physical principles (dashed line in **Figure S2E**, formula in methods and legend), which is comparable to the ∼ 10^11^ M^-1^s^-1^ measured experimentally for binding of single HIV-1 virions (sizes similar to pHEL) to target cells ^64^. Meanwhile, observed activation rates (*k*_obs_ values) were slower than the diffusion limit at high particle concentrations (1, 10 pM). These observations suggested a minimal kinetic scheme of a particle binding step followed by a cell-intrinsic activation step (**Figure S2E** inset), where particle binding is rate-limiting at low pHEL concentrations, while activation rate becomes limiting for high doses.

We next considered distinct models of B cell activation, where 1, 2, or 3 particle binding steps are required before a B cell can undergo a final cell activation step (**Figure S2F** and methods). We tested these models using maximum likelihood across experimental replicates and particle concentrations. We found that these models predicted distinct CDF curves at low (0.01 and 0.1 pM) particle concentrations where diffusion is rate-limiting, and could therefore be distinguished in those conditions based on the quality of fit to the experimental data (**Figure S2G**), whereas the differences among these models diminish at higher pHEL concentrations when the cell activation step was rate-limiting (**Figure S2H**). Notably, for diffusion limited conditions (0.1 and 0.01 pM pHEL), models where only one or two particle hits are required to activate a cell fit the data better than the 3-hit model (**Figure S2J-K**). This result was insensitive to the specific fluorescence threshold we chose to identify B cell activation (**Figure S2C, I-K**). This analysis suggests that one or two particles with high epitope density may be sufficient to activate B cells in this experimental system, and raises the intriguing possibility that naïve B cells are optimized to sense a single viral particle. Future single-particle tracking studies with high spatial and temporal resolution should further resolve the kinetics of pHEL binding and the subsequent activation of B cells with greater precision ^64^.

### Biased NF-κB and NFAT nuclear translocation by soluble and particulate Ag

To understand how early signaling events triggered by sAg and pAg produce qualitatively distinct fates (i.e. tolerance or immunity), we sought to examine transcriptional programming (**Figure 3A**). The balance between nuclear NFAT and AP-1 can regulate T cell tolerance; Ca^2+^-dependent nuclear NFAT translocation in the absence of AP-1 drives a transcriptional program comprised of negative regulators of TCR signal transduction ^30,65^. Similarly, anergic B cells exhibit constitutive nuclear NFAT localization and express an overlapping set of negative regulators, suggesting an analogous mechanism at play to limit B cell responses to signal 1 in the absence of signal 2 ^32,35,66^. Conversely, co-stimulation of B cells is essential for robust NF-κB activation by sAg. We previously showed that pAg produces maximal nuclear RelA/p65 translocation, while sAg required co-stimulation with CD40 ligation to do so ^49^.

**Figure 3:**
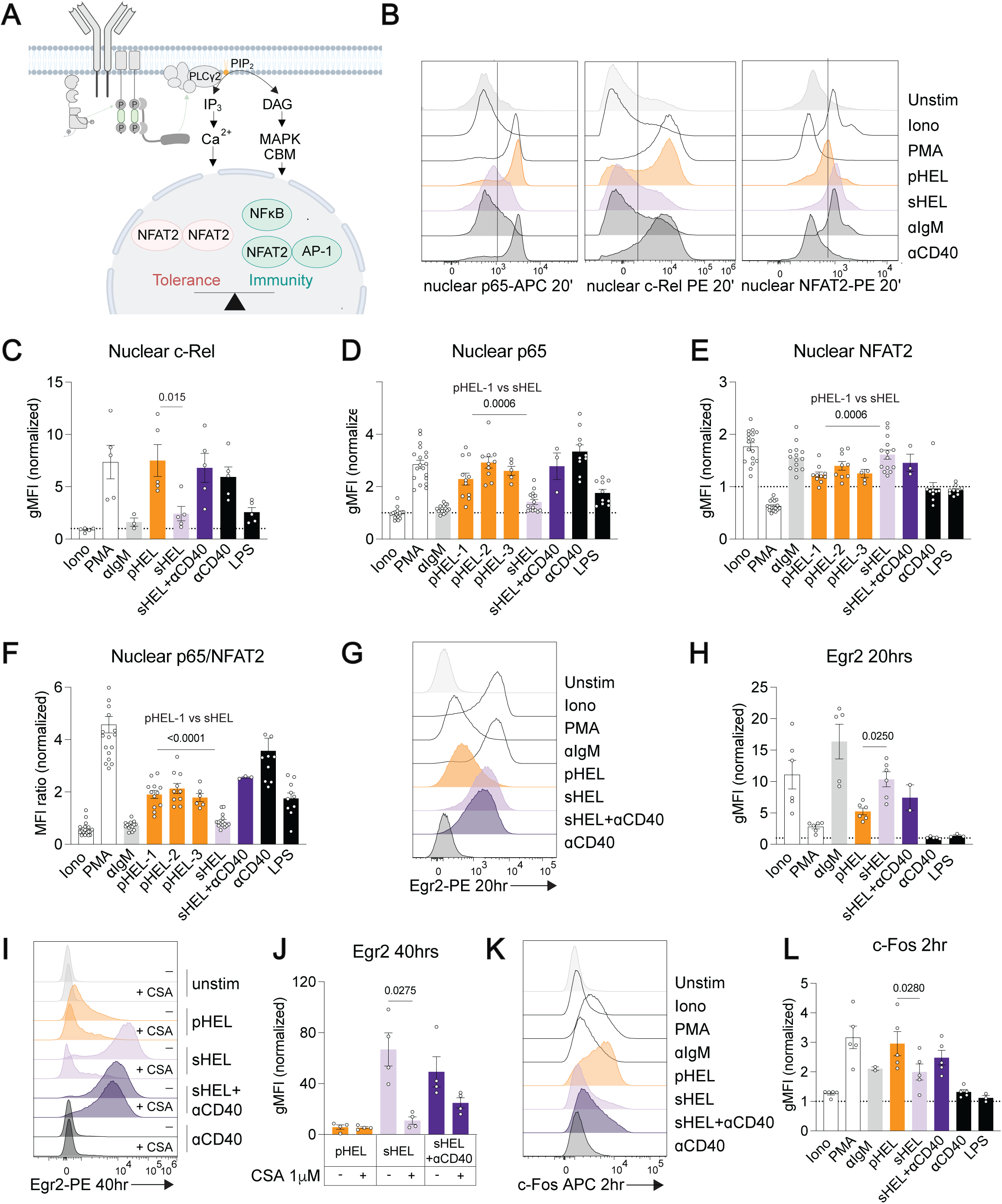
Biased NF-κB and NFAT nuclear translocation by soluble and particulate Ag 3. **A:** Proximal BCR signal transduction is translated to transcriptional fate via the products of PIP_2_ hydrolysis by PLCγ2 in the signalosome. IP_3_ generates cytosolic Ca^2+^ flux, resulting in the translocation of NFAT into the nucleus. DAG promotes MAPK and CBM (CARD11, BCL10, MALT1) complex activation, which support translocation of NF-κB and AP-1 transcription factor family members. In lymphocytes, NFAT homodimers promote a tolerogenic fate while NFAT-AP-1 heterodimers and NF-κB support an immune response. **3B:** B cells were isolated using negative selection from pooled MD4 splenocytes and lymph node cells and stimulated with the following: ionomycin 1μM, PMA 20ng/mL, anti-IgM 10μg/mL, pHEL 1pM, sHEL 1μg/mL, anti-CD40 100ng/mL or LPS 10μg/mL. After 20 minutes of stimulation, nuclei were isolated and stained for c-Rel, p65/RelA and NFAT2. NFAT2 and p65 staining was performed in tandem. Histograms depict flow cytometry data and are representative of >10 experiments. **3C-E:** Quantification of nuclear c-Rel (**C**), p65 (**D**) and NFAT2 (**E**) data shown in **3B**. Geometric mean fluorescence intensity (gMFI) was normalized to unstimulated conditions for each experiment. A dashed line at 1 represents baseline unstimulated MFI. Each dot represents an individual experimental replicate. p values were acquired by performing paired Student’s t-test. **3F:** Ratio of normalized MFIs of nuclear p65 to NFAT2. Each data point represents the ratio of p65 to NFAT2 from an individual experiment. **3G:** Pooled MD4 splenocytes and lymph node cells were stimulated overnight with stimuli at doses as denoted in **3A**, followed by intracellular staining for Egr2. Histograms are representative of 5 independent experiments. **3H:** Quantification of data in **3G**. gMFI values were normalized to unstimulated controls. p values were acquired by performing paired Student’s t-test. **3I:** Pooled MD4 splenocytes and lymph node cells were pre-incubated with Cyclosporin A for 30 minutes prior to stimulation for 40 hours and intracellular staining for Egr2. Histograms are representative of three independent experiments. **3J:** Quantification of data in **3I**. gMFI values were normalized to unstimulated controls. p values were acquired by performing paired Student’s t-test. **3K:** Pooled MD4 splenocytes and lymph node cells were stimulated for 2 hours and stained intracellularly for c-Fos. Histograms are representative of five independent experiments. **3L:** Quantification of data in **3K**. gMFI values were normalized to unstimulated controls. p values were acquired by performing paired Student’s t-test.

We examined nuclear translocation of these key transcription factors by staining B cell nuclei isolated post-stimulation to detect RelA/p65 and NF-κB c-Rel subunits and NFAT2, the predominant NFAT isoform expressed in B cells (**Figure S3A, B**). Relative to CD4+ T cells, B cells induce higher maximal nuclear NFAT2 (but lower NFAT1) with ionomycin and higher nuclear p65 in response to the DAG-mimetic PMA (**Figure 3SC, D).**

We observed robust, near-maximal nuclear RelA/p65 and c-Rel in response to pAg, comparable to that induced in response to PMA or CD40 stimulation (**Figure 3B-D**). By contrast, sAg or anti-IgM each induced only modest NF-κB activation, and this was enhanced by co-stimulation through CD40. Of note, we previously demonstrated that activation of MAPK and NF-κB pathways by pHEL was independent of Myd88 and IRAK1/4 signaling ^49^.

Surprisingly, although pAg triggers robust and prolonged Ca^2+^ mobilization (**Figure 1I**, **Figure 2A**), this stimulus does not induce maximal NFAT2 nuclear translocation (**Figure 3B, E**). Rather, pAg produces consistently lower NFAT2 translocation than anti-IgM and sAg. These soluble stimuli, by contrast, produce maximal NFAT2 translocation, comparable to that seen with high dose ionomycin. Both NFAT1 and NFAT2 nuclear translocation exhibit a similar pattern of response to these stimuli (**Figure S3B**). Critically, key controls validate NFAT nuclear staining, including induction with ionomycin, and abrogation of nuclear NFAT with either EGTA chelation of extracellular Ca^2+^ or treatment with the calcineurin inhibitor Cyclosporin A (**Figure S3E-H**).

Because NF-κB mediates immunogenic B cell responses, while nuclear NFAT alone is associated with tolerance in lymphocytes, we reasoned that the relative nuclear concentration of these two transcription factors might program B cells for different fates. As our NFAT2 and RelA/p65 nuclear staining was performed simultaneously, we were able to calculate the ratio of these transcription factors on a single cell level (**Figure 3F**). We found, as expected, that PMA, anti-CD40, and LPS produced robust NF-κB activation without inducible NFAT translocation, consistent with absence of cytosolic Ca2+ entry. Strikingly, pAg consistently favors NF-κB over NFAT translocation, resembling the ratio produced by LPS. By contrast, both anti-IgM and sAg stimulation resemble ionomycin, greatly favoring NFAT over NF-κB. Addition of signal 2 in the form of CD40 shifted this ratio to favor NF-κB. Importantly, we observed this trend across multiple, independently synthesized high density pHEL particles (pHEL-1, 2, 3), suggesting that this virus-like structure reproducibly triggers a biased pattern of nuclear transcription factor translocation **(Figure 3F).**

To further validate these observations, we profiled expression of a transcriptional target of NFAT in the absence of AP-1. Egr2 is a Ca^2+^-dependent primary response gene for whose transcription NFAT is necessary and sufficient, and which plays an important role in T and B cell anergy ^31,67^. We found, as expected, that ionomycin but not PMA were sufficient to induce intracellular Egr2 (**Figure 3G, H**), while the calcineurin inhibitor Cyclosporin A completely abrogated its induction (**Figure 3I, J**). While Egr2 is robustly induced by sAg and anti-IgM, it is minimally upregulated by pAg (**Figure 3G-J**). This suggests that nuclear NFAT following pAg stimulation evades an NFAT-only tolerogenic program and instead forms NFAT:AP1 heterodimers to drive an immunogenic program. Indeed, pAg robustly induces c-Fos protein expression, a component of AP-1, while sAg does not (**Figure 3K, L**). We confirmed that c-Fos induction is MAPK-dependent (**Figure S3I**), consistent with earlier results demonstrating maximal pErk induction by pAg but not sAg (**Figure 1D**)^49^. As a proxy for determining the relative abundance of NFAT:AP-1 activating dimers to NFAT:NFAT homodimers, we plotted the ratio of c-Fos to nuclear NFAT2 across stimuli (**Figure S3J**). pAg promotes more relative c-Fos than NFAT2, and a similar trend was observed with pErk (**Figure S3K**).

### DAG, but not calcium, is limiting for NF-κB activation by soluble antigen

pAg induces robust MAPK activation (**Figure 1D)** and Ca^2+^ mobilization (**Figure 1I**, **Figure 2A**), signaling events dependent on the second messengers diacylglycerol (DAG) and inositol-trisphosphate (IP3), respectively **(Figure S4A)**. These molecules are generated by PLCγ2-mediated hydrolysis of PI(4,5)P_2_, a process we previously showed was amplified by pAg ^17,49^. BCR signaling is linked to NF-κB activation by classical PKC which requires both Ca^2+^ and DAG for activation ^68^. Importantly, we show that NF-κB activation by pAg, sAg, and anti-IgM (but not CD40) is PKC-dependent and calcium-dependent (**Figure S4B-E**). This shows that pAg robustly activates NF-κB through the canonical BCR signaling pathway.

We sought to determine which of these signaling arms, DAG or IP3/Ca^2+^, drive robust p65 induction by pAg, and conversely, which arm was limiting for NF-κB activation by soluble BCR stimuli. To do so, we stimulated MD4 B cells with anti-IgM, sAg or pAg in the presence of varying doses of the DAG-mimetic PMA or the Ca^2+^ ionophore ionomycin. We selected concentrations of these reagents across a broad range to capture sub-maximal doses that could reveal synergy with sAg stimuli. We then isolated B cell nuclei to detect translocation of RelA/p65 **(Figure 4A)**. PMA at a saturating dose (20ng/mL) led to maximal nuclear p65 across all stimulation conditions as expected. Interestingly, at an intermediate, submaximal dose (2ng/mL) that is insufficient to induce robust NF-κB on its own, PMA synergizes with ionomycin, anti-IgM, and to some extent with sHEL to promote maximal p65 **(Figure 4B)**. By contrast, even a high dose of ionomycin sufficient to produce maximal nuclear NFAT translocation failed to boost p65 activation in response to anti-IgM or sHEL **(Figure 4A, S4F, G)**. These data suggest that DAG but not Ca^2+^ are limiting for NF-κB activation by sAg.

**Figure 4:**
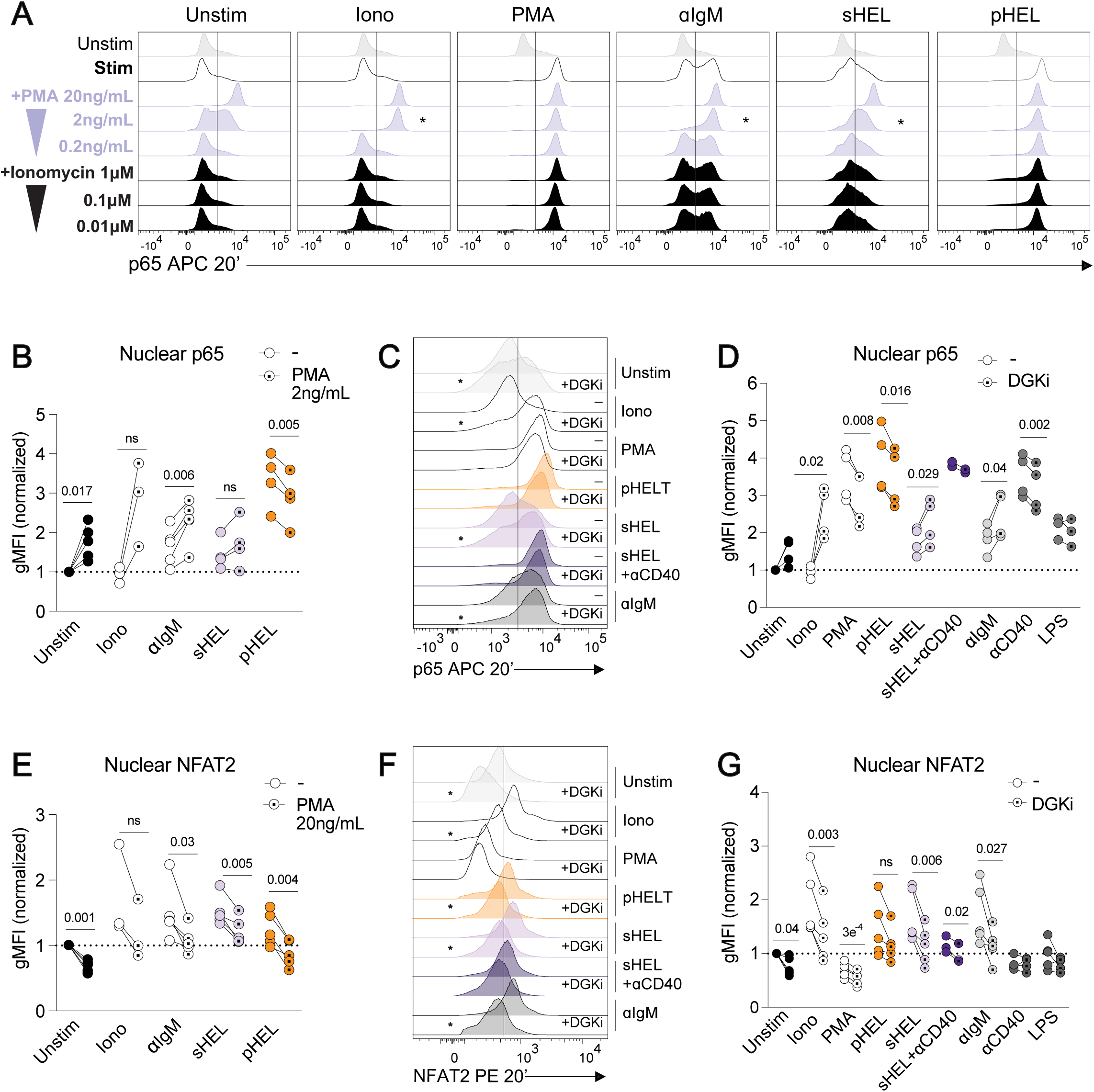
DAG, but not calcium, is limiting for NF-κB activation by soluble antigen 4. **A:** p65 staining in B cell nuclei. MD4 B cells were isolated from pooled splenocytes and lymph node cells and stimulated for 20 minutes with PMA (20ng/mL), ionomycin (1µM), anti-IgM (10μg/mL), sHEL (1μg/mL) or pHEL (1pM) alone or with additional titrations of PMA (20, 2, 0.2ng/mL) or ionomycin (1, 0.1, 0.01µM). Data representative of five independent experiments. **4B:** Quantification of data in **4A** highlighting dual stimulation conditions with mid doses of PMA (2ng/mL) or ionomycin (0.1µM). gMFI values were normalized to unstimulated controls. Each paired dataset represents an individual experimental replicate. P values acquired via Student’s paired t-test. **4C:** p65 staining in B cell nuclei. B cells were incubated with DGK inhibitor (R59949 100µM) for 30 minutes prior to stimulation for 20 minutes. Data representative of at least four independent experiments. **4D:** Quantification of data in **4C**. gMFI values were normalized to unstimulated controls. Each paired dataset replicates an individual experimental replicate. P values acquired via Student’s paired t-test. **4E:** Quantification of NFAT2 nuclear staining data in **S4I**. MD4 B cells were isolated from pooled splenocytes and lymph node cells and stimulated for 20 minutes with PMA (20ng/mL), ionomycin (1µM), anti-IgM (10μg/mL), sHEL (1μg/mL) or pHEL (1pM) alone or with additional titrations of PMA (20, 2, 0.2ng/mL) or ionomycin (1, 0.1, 0.01µM). gMFI values were normalized to unstimulated controls. Each paired dataset replicates an individual experimental replicate. P values acquired via Student’s paired t-test. **4F:** NFAT2 staining in B cell nuclei. B cells were incubated with DGK inhibitor (R59949 100µM) for 30 minutes prior to stimulation for 20 minutes. **4G:** Quantification of data in Figure 4F. gMFI values were normalized to unstimulated controls. Each paired dataset replicates an individual experimental replicate. P values acquired by performing paired Student’s t-test.

We sought a complementary strategy to test whether DAG is limiting for NF-κB activation downstream of sAg stimuli. Endogenous DAG is metabolized to phosphatidic acid (PA) by diacylglycerol kinase (DGK) **(Figure S4A)** ^69^. Therefore, we utilized a well-characterized small molecule inhibitor of DGK (R59949, DGKi) to boost endogenous DAG concentration by blocking its turnover ^70^. We confirmed that DGKi markedly potentiates pErk induction by anti-IgM (**Figure S4H**). We observed that inhibition of DGK boosted nuclear p65 in response to ionomycin, anti-IgM and sAg (**Figure 4C, D**). DGKi is bioactive in both T and B cells, sufficient to synergize with ionomycin to activate NF-κB (**Figure S4H**). Nevertheless, DGKi synergizes with basal signaling only in B cells, but not T cells, suggesting enhanced basal signaling in the former (**Figure S4I**). Collectively, these results suggest that DAG is limiting for NF-κB activation by sAg, and that DGKs play an important role in constraining B cell responses to signal 1 in the absence of signal 2.

### Nuclear NFAT is suppressed by DAG

We were surprised to observe modest nuclear NFAT2 following pAg stimulation despite remarkably enhanced and prolonged cytosolic Ca^2+^ responses (**Figures 1I, 2A, 3B, 3E**). Because this muted NFAT response in turn resulted in limited Egr2 induction, a marker of anergic NFAT signaling (**Figure 3G-J**), we next sought to understand the mechanism for uncoupling of Ca^2+^ and nuclear NFAT. We consistently observed “suppression” of nuclear NFAT2 in B cells below basal levels upon treatment with PMA (**Figure 3B, E**). We also find that PMA suppresses nuclear NFAT2 in cells stimulated through the BCR with both soluble and particulate stimuli. This is accomplished in a dose-dependent manner; across conditions, PMA suppresses NFAT2 at high (20 ng/mL) and intermediate (2 ng/mL) doses (**Figure 4E, S4J, K**). Importantly, boosting endogenous DAG with DGKi also suppresses nuclear NFAT2 across all stimuli and even in unstimulated B cells (**Figure 4F, G, S4L**). This effect is most appreciable in cells stimulated with strong inducers of NFAT2: ionomycin, anti-IgM, and sAg.

PMA-mediated suppression of NFAT2 more than NFAT1 is evident in B cells (**Figures S3B-D**). Moreover, suppression of nuclear NFAT2 appears to be largely specific to B cells and is less apparent in T cells following either PMA or DKGi treatment (**Figure S3C, D, S4L**). It is possible this difference reflects higher basal signaling and nuclear NFAT2 in B cells than in T cells (**Figure S3C, D**).

Collectively, these data support a model in which robust DAG production triggered by pAg not only synergizes with Ca^2+^ to maximally activate NF-κB but also suppresses nuclear NFAT. Conversely, cytosolic Ca^2+^ produced by soluble Ag stimuli like sHEL and anti-IgM is sufficient to drive nuclear NFAT translocation, but concomitant modest DAG production is limiting for NF-κB and insufficient to suppress nuclear NFAT. This is consistent with a low Ca^2+^ threshold for nuclear NFAT localization described in lymphocytes ^32,35,71,72^. Together this wiring enables B cells to modulate the relative nuclear localization of NFAT and NF-κB in response to soluble or particulate Ag, rather than simply titrating total amplitude of the response.

### Soluble and particulate antigen induce distinct transcriptional landscapes

Given the distinct nuclear translocation of NFAT and NF-κB triggered by soluble and particulate Ag, we postulated that the transcriptional landscape of B cells activated by distinct formats of antigen might diverge profoundly. To test this hypothesis, we stimulated mature follicular MD4 B cells isolated from lymph node for 16 hours with either pHEL, sHEL alone, sHEL with anti-CD40 stimulation to mimic T cell help, or with LPS to model a classic mitogenic “danger signal” and then performed RNAseq **(Figure 5A**, **Table 1a)**. Our goal was to identify *qualitatively* distinct transcriptional modules induced by these conditions which represent classic T-dependent and T-independent stimuli. In particular, we sought to gauge whether pHEL stimulation mimicked signal 1 delivered together with signal 2 (sHEL + anti-CD40) or rather resembled a classic “stand-alone” mitogenic danger signal such as LPS. We selected equipotent doses of pHEL and sHEL that induce comparable induction of the activation marker CD69 and confirm this is independent of IRAK1/4 (**Figure 5B-C, S5A**). We selected a robust dose of CD40 co-stimulation sufficient to drive maximal NF-κB activation (**Figure 3B-D**).

**Figure 5:**
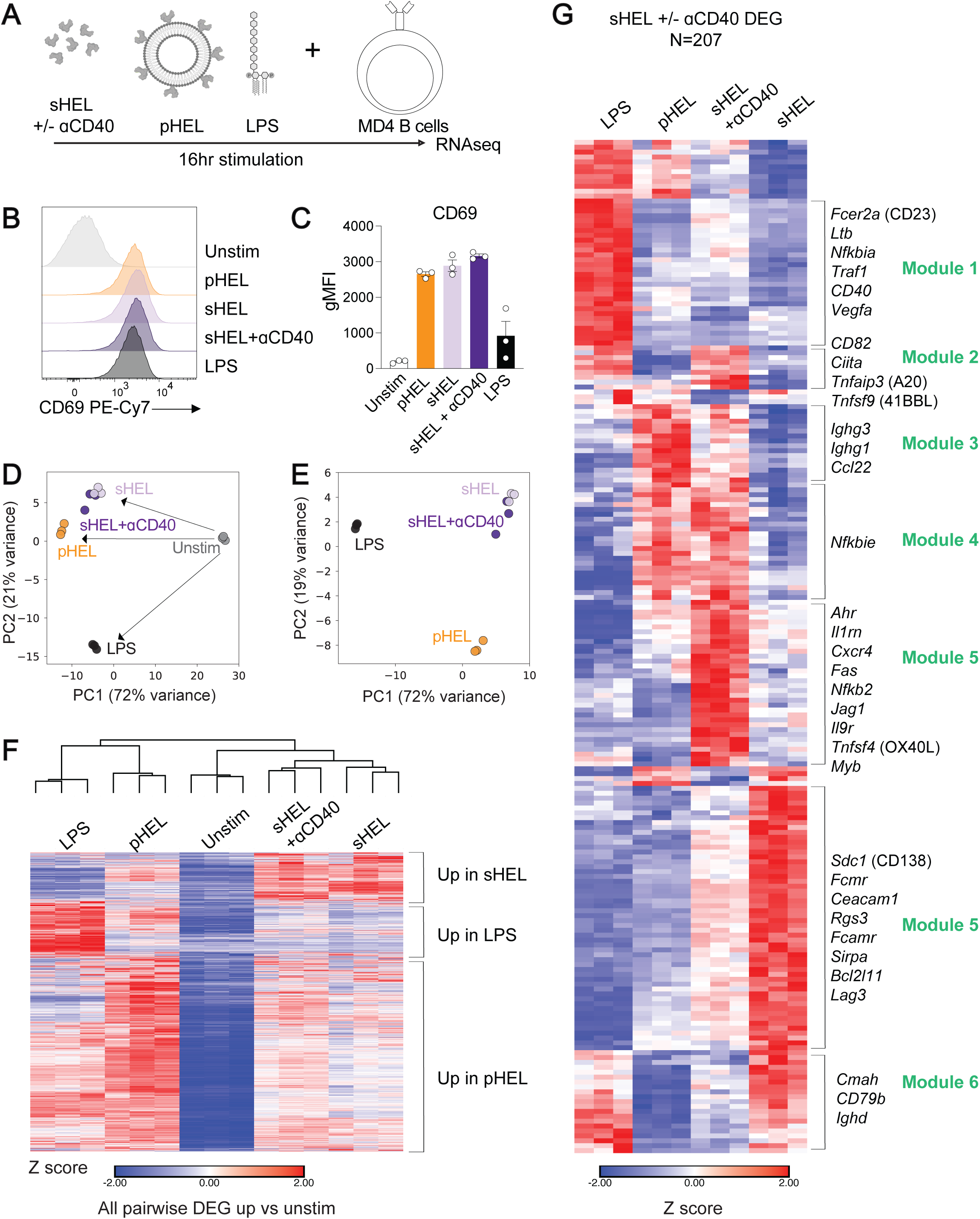
Soluble and particulate antigen induce distinct transcriptional landscapes 5. **A:** Experimental design for RNAseq. B cells were isolated from MD4 lymph nodes from three mice and stimulated in biological triplicate for 16 hours with sHEL (1μg/mL), sHEL+αCD40 (100ng/mL), pHEL (1pM) or LPS (10μg/mL) prior to harvest for sequencing. **5B:** CD69 surface staining of stimulated cells at 16hr timepoint. Histograms representative of additional biological replicates. **5C:** Quantification of data in **5B**. **5D:** Principal component analysis (PCA) plot of all experimental conditions. RNAseq data was analyzed using DESeq2 platform. **5E:** PCA plot of all stimulation conditions. **5F:** Heatmap of all pairwise differentially expressed genes (DEGs) that are up in stimulated cells versus unstimulated cells (padj<0.05, |log2FC|>=1). Each column represents an individual biological sample. **5G:** Heatmap of all differentially expressed genes (DEGs) between sHEL-stimulated cells and cells stimulated with sHEL+αCD40 (N=207, padj<0.05, |Log2FC|>0.5). Brackets highlight gene modules that are upregulated in distinct patterns between stimulation conditions.

Principle component analysis (PCA) suggests that all stimulation conditions diverge from unstimulated controls, and this accounts for the majority of the variance among the sample transcriptomes **(Figure 5D)**. PCA on stimulated samples only revealed three broad sample clusters corresponding to LPS, pHEL, and sHEL, with most profound variance driven by LPS (**Figure 5E**). Surprisingly, CD40 co-stimulation of sHEL-activated B cells accounted for relatively limited variance. Importantly, pHEL does not cluster with this group nor with LPS, suggesting that the transcriptional signature of pAg-stimulated cells diverges markedly from CD40 co-stimulation and does not simply recapitulate a canonical danger signal.

Next, we generated a heatmap of all pairwise differentially expressed genes (DEGs) across our samples that were upregulated relative to unstimulated B cells ((N=3942, padj<=0.05, |log2FC|>=1); **Figure 5F**). Consistent with global PCA, hierarchical clustering analysis of pairwise DEGs reveals high similarity among sHEL samples +/− CD40 stimulation and interestingly reveals similarities between LPS and pHEL stimulated samples. We observed several distinct gene expression modules; a group of transcripts most highly expressed in sHEL-stimulated samples +/− CD40, a module of LPS-induced transcripts, and a surprisingly large set of pHEL-induced transcripts, a subset of which are shared with LPS (**Figure 5F**).

To understand this transcriptional divergence at a more granular level, we performed a *separate* PCA to identify the transcripts whose differential expression contribute highly to the top four principal components and thus help account for key variance among groups (**Figure S5B, C, Table 1b**). This identified genes whose expression was differentially regulated by LPS (PC1), others specific to pHEL (PC2), and finally a set that distinguished CD40 co-stimulation (PC4). Among the genes that drive PC1 segregating sHEL samples was *Egr2* (**Figure S5B, C**), recapitulating its protein expression pattern (**Figure 3G-J**).

### pAg induces a transcriptome that diverges from canonical signal 2

We initially hypothesized that pHEL might recapitulate the transcriptional program induced by the combination of signal 1 (sHEL) and signal 2 (CD40). Yet our global analyses revealed that addition of CD40 co-stimulation not only differed from pHEL, but accounted for a relatively small degree of variance (**Figure 5D-F, S5B** e.g. PC4). To further explore this, we identified a constrained set of genes that were differentially expressed between sHEL-activated B cells with or without CD40 stimulation (N=207, padj<0.05, |Log2FC|>0.5) (**Figure 5G**). Hierarchical clustering identified several discrete modules of co-regulated transcripts (**Figure 5F**).

We focused our attention first on genes induced by CD40 co-stimulation and identified modules that were shared by pHEL, including transcripts such as *Nfkbie* (IkBe), reflecting NF-κB activation feedback, and the Ccr4 ligand *Ccl22* which functions as a T cell chemoattractant ^73^. We also identified *Ighg1* and *Ighg3* (germline transcripts, GLTs) even at this early time point more highly induced by pHEL than CD40 (**Figure 5G, S5C**). Indeed, C57BL/6 mice immunized with pHEL produce IgG1 and IgG3 class-switched antibody even in the absence of T cell help ^47^.

Another set of transcripts was induced by either CD40 co-stimulation or LPS (two distinct forms of “signal 2”), but not pHEL. These included other NF-κB regulated genes, including feedback regulators *Nfkbia* (IkB) and *Tnfaip3* (A20) (**Figure 5G**). Although we observe robust induction of both RelA/p65 and c-Rel by pHEL (**Figure 3B-D**), the kinetics and heterodimers formed by the NF-κB subunits may differ among co-stimulatory inputs to produce distinct NF-κB modules ^74–76^. Importantly, this data set also identifies known CD40-regulated transcripts including *Fcer2a* (CD23) as well as those uniquely regulated by CD40 (*Fas*, *Nfkb2* (p100), *Tnfsf4* (OX40L), and *Il9r*) (**Figure 5G, S5C;** GSE159854)^77–79^.

Strikingly, we also identified a module of genes that is highly expressed in sHEL-stimulated B cells but down-regulated with CD40 co-stimulation and minimally expressed with pHEL or LPS-activation (**Figure 5G**). These include negative regulators of B cell signaling such as inhibitory co-receptors *Ceacam1*, *Sirpa*, and *Lag3* as well as the pro-apoptotic factor *Bcl2l11 (*BIM) and additional markers previously associated with B cell anergy such as *Sdc1* (CD138)^66^.

### Soluble, but not particulate, antigen induces an anergic transcriptome, which is only partially alleviated by the addition of signal 2

It has long been argued that signal 1 in the absence of signal 2 results in B cell anergy. Indeed, several genes implicated in both B cell and T cell anergy were induced by sHEL but not be pHEL or LPS, suggesting that sHEL induces an anergy-associated transcriptome that is evaded by pHEL (**Figure 5F**). These include *Egr2/*3 and the *Nr4a* transcription factor family members, primary response genes that are directly induced by NFAT in the absence of AP-1 (i.e. signal 1 without signal 2) ^80,81^, as well as the Nr4a-dependent gene *Eno3* (**Figure 5F, S5C, S6A**) ^82^. *Spry2* encodes a negative feedback regulator of MAPK signaling that re-enforces nuclear NFAT without AP-1 ^83^. Strikingly, *Dgka* and *Dgkz* – well-described genes that contribute to T cell anergy by restricting DAG and downstream MAPK/NF-κB activation – are also more robustly upregulated by sAg than by pAg **(Figure S6B)** ^69,84,85^. DGKs constrain B cell responses as well ^86^; indeed, we showed that DAG tuning regulates the balance between tolerogenic and immunogenic TF signaling (**Figure 4**).

To further investigate the transcriptional program induced by sHEL, we compared our dataset to a bona fide anergic transcriptome identified in the well-studied HEL-mouse model. In this model, IgHEL B cells develop in the context of model self-antigen sHEL (IgHEL Tg x sHEL Tg; MD4/ML5 line), and acquire many canonical features of B cell anergy, including IgM downregulation, functional and biochemical unresponsiveness, and a unique transcriptome ^4,23,50,66^. This anergy-associated signature was highly enriched among sHEL-activated B cells in our dataset relative to pHEL (**Figure 6A**, **B** normalized enrichment score (NES)=1.99 sHEL vs pHEL, **Table 1c, d**)^87^. Interestingly, this anergic transcriptome was minimally impacted by CD40 co-stimulation (**Figure 6B**, NES=1.39 sHEL + anti-CD40 vs pHEL). In other words, CD40 is insufficient to suppress this program while pHEL uniquely evades it.

**Figure 6:**
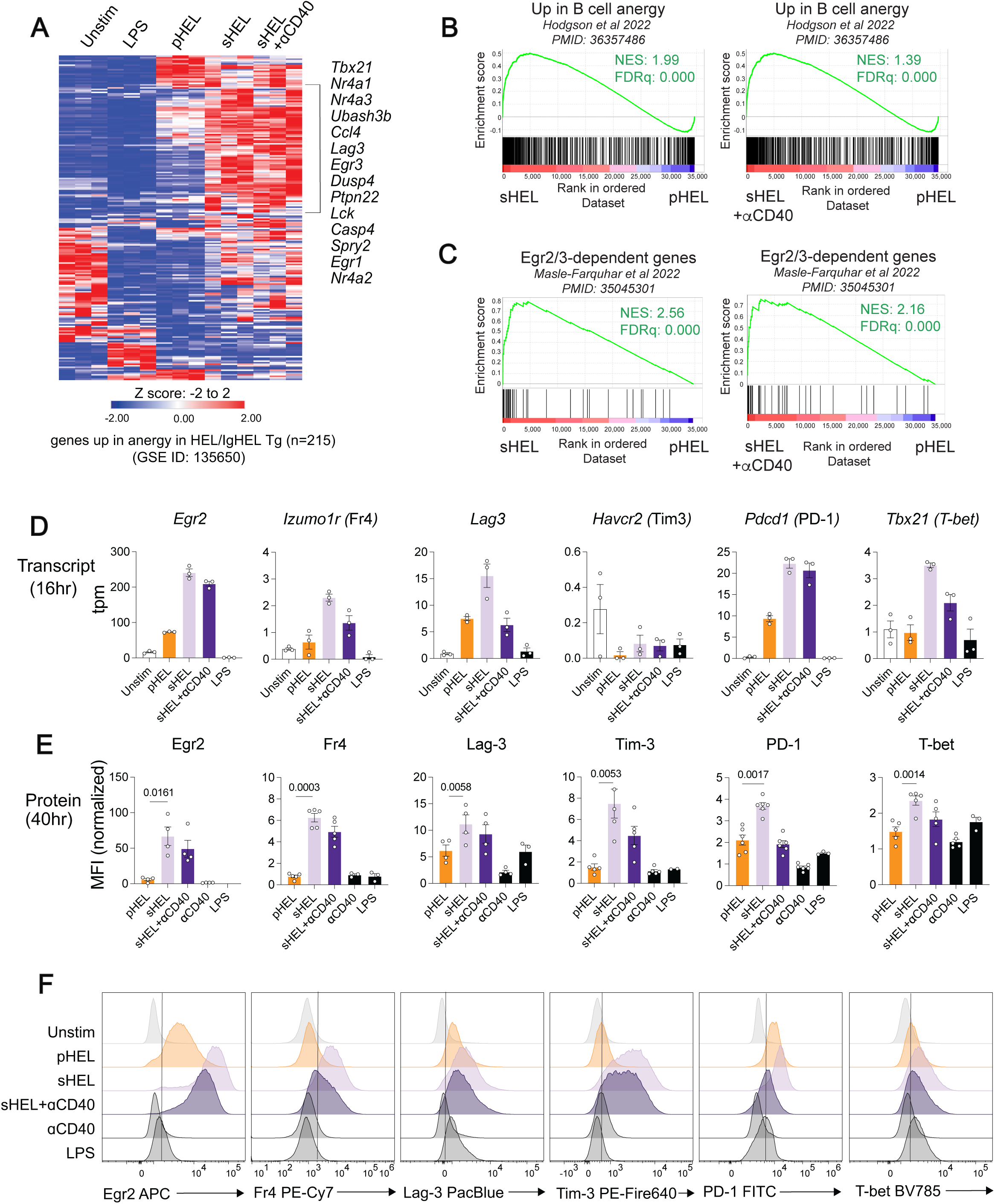
Soluble, but not particulate, antigen induces an anergic transcriptome, which is only partially alleviated by the addition of signal two 6. **A:** Heatmap depicts our RNAseq expression data across the set of genes upregulated in IgHEL-sHEL anergic B cells ^87^. Annotation highlights anergy-associated genes enriched in sHEL- and sHEL+aCD40-stimulated B cells. **6B:** GSEA performed to compare genes upregulated in sHEL- (left) or sHEL+aCD40-(right) versus pHEL-stimulated B cells to the data set of genes upregulated in ex vivo anergic B cells versus naïve B cells (GSE135650). NES = normalized enrichment score. FDRq = false discovery rate q-value. **6C:** GSEA performed to compare genes upregulated in sHEL- (left) or sHEL+aCD40-(right) versus pHEL-stimulated B cells to the data set of genes upregulated in *Egr2−/−Egr3−/−* versus wildtype follicular B cells (ENA PRJEB48165). NES = normalized enrichment score. FDRq = false discovery rate q-value. ^31^. **6D:** Quantification of anergy-associated transcripts from RNAseq data. **6E:** Quantification of anergy-associated protein expression at 40 hours. Pooled splenocytes and lymph node cells from MD4 mice were cultured for 40 hours with pHEL (1pM), sHEL (1μg/mL), sHEL+αCD40, αCD40 (100ng/mL) or LPS (10μg/mL) and then stained for surface and intracellular anergy markers. gMFIs were normalized to unstimulated controls. Each data point represents an experimental replicate. Data representative of at least four independent experiments. P values acquired from Student’s paired t-test. **6F:** Representative flow cytometry data from that described in **6E**.

NFAT in the absence of AP-1 is a key driver of B cell anergy. In B cells, it was recently shown that the NFAT-dependent genes *Egr2* and *Egr3* contribute functionally to the anergy program in the MD4/ML5 model ^31^. Egr2/3-dependent genes were also enriched in sHEL-activated B cells relative to pHEL stimulation (**Figure 6C**, NES=2.56). Here again, signal 2 in the form of CD40 co-stimulation did not reverse this Egr2/3-dependent program (NES=2.16).

Of note, several inhibitory co-receptors were among the genes induced by sAg, including *Havcr1* (Tim1), *Havcr2* (Tim3), *Lag3*, and *Pdcd1* (PD-1), several of which we confirmed at the protein level (**Fig S6A, 6D-F**). We additionally validated several other key anergy-associated genes including *Egr2* (**Figure 3, Figure S5C**), the T cell anergy marker Fr4 (*Izumo1r*), and the atypical memory B cell marker T-bet *(Tbx21)* (**Figure 6D-F)**. Many of these surface markers were detectable only at later time points of stimulation, trailing the transcriptional program. The addition of CD40 co-stimulation only partially suppressed transcript and protein expression induced by sHEL. By contrast, pHEL induced modest expression of some of these proteins, and in some cases evaded induction entirely (as for Egr2, Fr4, and Tim3). The expression of these markers was partially dependent on NFAT; addition of the calcineurin inhibitor Cyclosporin A (CSA) reduced Egr2, T-bet and PD-1 induction in response to sHEL stimulation (**Figure 3I-J, Figure S6C-D**).

Thus, sAg stimulation rapidly establishes an anergy-like transcriptome within B cells after just 16hrs and additional T cell help in the form of CD40 co-stimulation may only partially rescue this phenotype. In contrast, pAg appears to evade this “tolerance” program almost entirely.

### Particulate but not soluble Ag induces a Myc-dependent transcriptional program

Our global heatmap revealed a broad module of genes that were uniquely induced by pHEL stimulation (**Figure 5F**). Many of these pHEL-induced transcripts were involved in cell cycle (*Cdk, Cdc* genes), DNA replication (*Mcm* genes, Polymerase subunits), RNA processing (*Hnrnp*s), and translation (*Eif, Rpl, Rps* genes), a program that is orchestrated by c-Myc ^88–90^. We identified a very strong enrichment for Myc-hallmark genes in pHEL samples (**Figure 7A,B** NES = 4.41 pHEL vs sHEL). This signature was enhanced by CD40 co-stimulation (**Figure S7A**, NES = 2.59 sHEL+anti-CD40 vs sHEL), but was much more highly induced by pHEL (**Figure 7A,C** NES=3.91 pHEL vs sHEL+aCD40). Notably, LPS, strong mitogenic stimulus for B cells, also induces a Myc-hallmark program, but pHEL more robustly drives this program despite a vast difference in dose (**Figure 7A**; 10μg/ml or ∼1μM LPS vs. 1pM pHEL). We independently confirmed that the Myc-dependent transcriptome triggered downstream of LPS in B cells was also enriched in pHEL-stimulated samples (**Figure S7B, C**)^88^.

**Figure 7:**
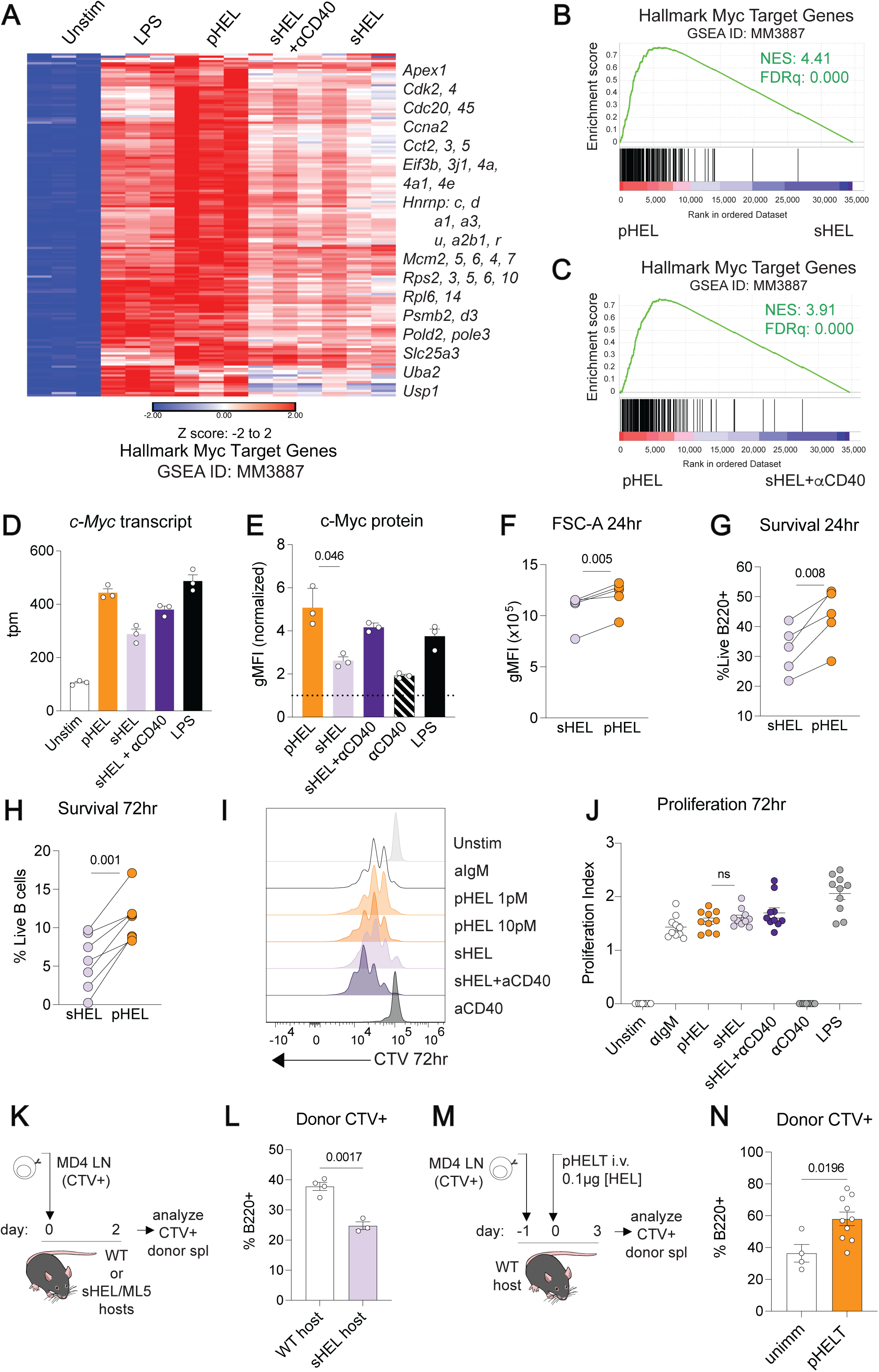
Particulate but not soluble Ag induces a Myc-dependent program 7. **A:** Overlay of RNAseq data with hallmark Myc-target genes (GSEAID: MM3887). **7B-C:** GSEA performed to compare genes upregulated in pHEL- versus sHEL- **(B)** or sHEL+aCD40- **(C)** stimulated B cells to the data set of hallmark Myc target genes (GSEAID: MM3887). NES = normalized enrichment score. FDRq = false discovery rate q-value. **7D:** Quantification of c-*Myc* transcript from RNAseq data. **7E:** c-Myc protein expression in B cells following overnight culture with various stimuli. Data represent gMFIs normalized to unstimulated controls. **7F:** FSC-A raw gMFIs of sHEL- and pHEL-stimulated cells overnight. P values acquired from Student’s paired t-test. **7G-H:** Viability (% live B220+) of sHEL- and pHEL-stimulated cells at 24 **(G)** or 72 **(H)** hrs, in the absence of BAFF. P values acquired from Student’s paired t-test. **7I:** MD4 lymphocytes were loaded with Cell Trace Violet (CTV) vital dye and cultured with denoted stimuli for 72hrs. Representative histograms depict CTV dilution in live B220+ cells. Data representative of ten biological replicates. **7J:** Quantification of proliferation indices, using vital dye dilution as show in **7I**. P values acquired from Student’s paired t-test. **7K:** MD4 lymph node cells were labeled with CTV and transferred into C57BL/6 wildtype or sHEL-expressing ML5 hosts. After two days, spleens were harvested from host animals and CTV+ donor cells were analyzed by flow cytometry. **7L:** Quantification of data from experiment outlined in **7K**, showing percentage of B220+ cells within CTV+ donor cells in WT or sHEL hosts. Each data point represents a biological replicate. P values acquired from Student’s paired t-test. **7M:** MD4 lymph node cells were labeled with CTV and transferred into C57BL/6 wildtype hosts one day before immunization with pHELT. Three days post immunization, spleens were harvested from host animals and CTV+ donor cells were analyzed by flow cytometry. **7N:** Quantification of data from experiment outlined in **7M** showing percentage of B220+ cells within CTV+ donor cells in WT hosts immunized with pHELD and unimmunized controls. Each data point represents a biological replicate. P values acquired from Student’s paired t-test.

Robust *c-Myc* transcript induction by pHEL corresponded well to c-Myc protein induction, similar to prior observations (**Figure 7D, E)** ^49^. Notably, CD40 alone was insufficient to drive c-Myc protein induction, requiring signal 1 (Ag) to do so **(Figure 7E).** We verified functional sequelae of c-Myc ^88,90^, observing increased cell size (forward scatter/FSC) and viability following stimulation with pHEL relative to sHEL **(Figure 7F, G)**.

### Particulate Ag partially rescues B cells from Ag-induced cell death and drives robust proliferation

Ag-activated B cells that fail to receive co-stimulation within a limited time window are consigned to anergy or apoptosis ^24,25^. Indeed, reduced survival is a characteristic phenotype of anergic B cells that receive chronic signal 1 without signal 2 ^91–94^. B cells stimulated with sAg are thus highly dependent on CD40 co-stimulation for survival (**Figure S7D**). By contrast, pAg even in the absence of co-stimulation promotes B cell survival (**Figure 7H**). However, B cells activated with either pHEL or sHEL remain highly dependent on BAFF for their survival, while CD40 or LPS can largely substitute for BAFF signaling by signaling to the non-canonical NF-κB pathway (**Figure S7E**) ^95,96^.

To understand what programs may account for this, we examined Bcl2 family member expression. LPS robustly drives specific pro-survival Bcl2 family transcripts (*Bcl2, Mcl1, Bcl2a1*), while CD40 promotes *Bcl2a1* and *Bcl2l1* (Bcl-XL) ^75^. Strikingly, sHEL induces pro-apoptotic BH3-only genes *Bcl2l11* (BIM), *Bmf*, *Pmaip1* (NOXA), and *Bad* (**Fig S7F, S7G**). Interestingly, pHEL evades BH3-family member induction, but fails to drive highly robust pro-survival Bcl2 family transcripts, perhaps accounting for why pHEL stimulation remains BAFF-dependent.

We previously showed that B cell proliferation scales with sAg affinity and dose under BAFF-replete conditions where survival is not limiting ^49^. pHEL decorated with low affinity epitopes was sufficient to drive comparable proliferation in vitro, and extensive proliferation in vivo ^49^. Here again, we found that low pM doses of pHEL produce proliferation comparable to that achieved with high concentrations of high affinity sHEL in vitro (**Figure 7I, J, S7H**).

Marked differences between B cell responses to sHEL and pHEL are even more apparent in vivo under physiological BAFF conditions. To model this, we adoptively transferred labeled lymphocytes from IgHEL BCR Tg (MD4) donors into either WT or sHEL Tg (ML5) hosts; sHEL-activated B cells rapidly die within days (**Figure 7K, L**), consistent with prior reports ^91–94^. By contrast, transferred donor cells from MD4 mice in hosts that subsequently receive pHELT immunization (low dose of low affinity, high epitope density particles with no additional adjuvant) exhibit robust survival and expansion (**Figure 7M, N**)^49^.

### Particulate antigen-specific cells efficiently present antigen to activate T cells

A critical feature of the B cell response to Ag harboring T cell epitopes is capture, processing, and presentation of such epitopes on MHCII. Although we show that pHEL drives immunogenic T-independent responses in vitro and in vivo ^47^, multivalent stimuli including viruses can drive T cell-B cell collaboration. We next sought to understand how particulate and soluble Ag may impact B cell function as a professional APC.

To test B cells presentation of particle-derived immunogens, we purified HELD protein fused with the Eα_52-68_ peptide, sHELD-Eα (**Figure S8A**). We additionally conjugated this protein to liposomes to generate pHELD-Eα. Presentation of Eα_52-68_ on C57BL/6 MHCII I-A^b^ molecules can be specifically detected using the YAe monoclonal antibody ^97^(**Figure 8A**). After four hours of stimulation with either sHEL-Eα or pHELD-Eα, we were able to detect surface MHCII-Eα on IgHEL-specific B cells (**Figure 8B**). Interestingly, YAe fluorescence intensity scales with dose of HEL-Eα antigen irrespective of Ag format; B cells stimulated with 0.01μg/mL of sHEL-Eα showed similar YAe fluorescence as B cells treated with 10pM pHELD-Eα (∼0.01 μg/mL HEL-Eα protein). This suggests highly efficient capture and presentation of both particulate and soluble Ag. By contrast, CD69 upregulation by pHEL-Eα is much more robust than the equivalent concentrations of sHEL-Eα (10pM pHEL vs. 0.01μg/mL sHEL); indeed 1μg/mL of sHEL-Eα is required to drive CD69 upregulation comparable to 10pM pHELD-Eα, reflecting a 1000-fold potency gap, consistent with our prior studies^49^1.

**Figure 8:**
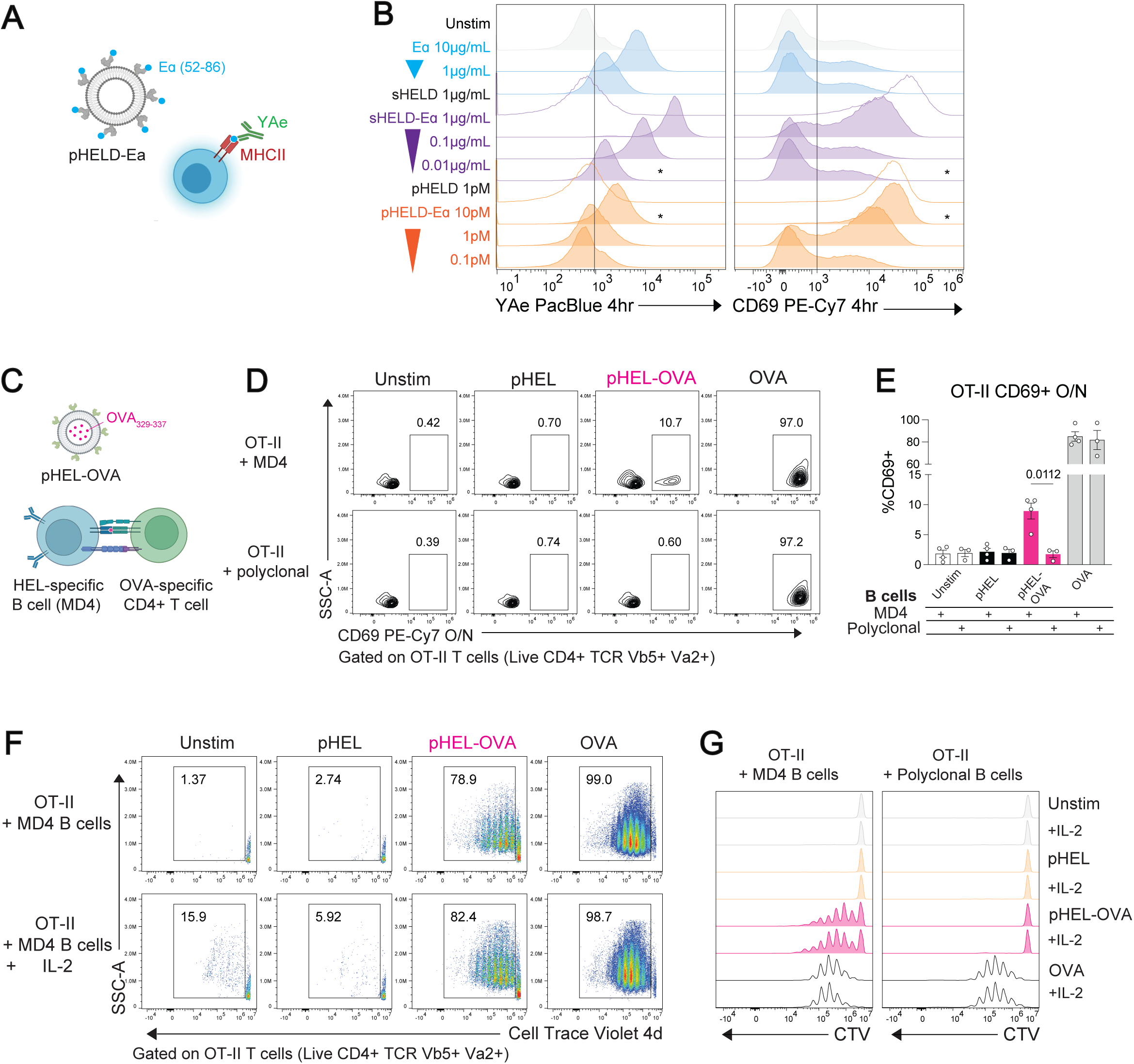
Particulate antigen-specific cells efficiently present antigen to activate T cells 8. **A:** Schematic for antigen presentation assay. pHEL particles were engineered to express the Eα peptide linked to HEL protein. The YAe antibody allows for direct recognition of the Eα peptide bound to C57BL/6 MHC II molecules. **8B:** Pooled MD4 splenocytes and lymph node cells were stimulated for 4hrs with Eα peptide alone, sHEL, ten-fold dilutions of sHE-Eα, pHELD, and ten-fold dilutions of pHELD-Eα. Histograms depict surface staining with YAe (left) and CD69 (right) on live B220+ cells. Stars denote matched HEL protein concentrations between sHEL-Eα and pHELD-Eα stimuli. Data representative of three experiments. **8C:** Schematic of B cell -T cell stimulation assay. MD4 B cells and OT-II CD4+ T cells were isolated from pooled spleen and lymph node cells from respective hosts. B and T cells were co-cultured and stimulated with pHEL particles encapsulating the OT-II specific OVA peptide (329-337). **8D:** OT-II CD4+ T cells were co-cultured with MD4 B cells or polyclonal B cells and noted stimuli (pHELT 10pM, pHELT-OVA 10pM, or OVA 5μM) overnight. Plots depict CD69 staining on OT-II T cells, denoted as live B220- Va2+ Vb5+ cells. Data representative of four experiments. **8E:** Quantification of data in **8D**. P values determined by Student’s paired t-test. **8F:** Isolated OT-II CD4+ T cells were loaded with CTV and co-cultured with MD4 B cells and denoted stimuli with or without additional IL-2 (5ng/mL). Plots depict CTV dilution in OT-IIs after four days. Data representative of three experiments. **8G:** Histograms displaying CTV dilutions by OT-II T cells with MD4 and polyclonal B cells after 96 hours of denoted stimulations. Data representative of three experiments.

While pMHCII-Eα on the surface of B cells scales with HEL-Eα Ag concentration, upregulation of CD86 is discordant and reflects particle potency (**Figure S8B**). This suggests that pAg delivers disproportionately robust T cell co-stimulation via CD86-CD28 engagement relative to pMHCII presentation.

Next, we evaluated whether pAg-stimulated B cells could serve as bona fide APCs to produce T cell activation. We generated pHEL particles containing encapsulated OVA_329-337_ (pHEL-OVA) and co-cultured CD4+ OT-II T cells with specificity for this epitope together with either HEL-specific MD4 B cells or polyclonal splenocytes (**Figure 8C**). We found that pHEL-OVA particles triggered CD69 upregulation in a subset of OT-IIs (**Figure 8D, 8E**). Critically, this activation was Ag-specific since it was not observed with polyclonal APCs, nor with pHEL particles lacking OVA peptide **(Figure 8D, E)**. Additionally, co-culture of MD4 B cells with pHEL-OVA and polyclonal CD4+ T cells resulted in no T cell activation **(Figure S8C)**. We also observed antigen-specific OTII proliferation in response to pHEL-OVA presented by MD4 B cells, even in the absence of exogenous IL-2 (**Figure 8F, G**). Thus, pAg-activated B cells, while not initially dependent on T cell help, can nevertheless serve as potent antigen-presenting cells and recruit T cells into an immune response.

## Discussion

It has been appreciated for decades that highly multivalent antigens can induce potent, T-independent B cell responses. This has long been attributed to extensive BCR cross-linking. However, we recently showed, using a new generation of pAg with defined molecular composition, that evasion of Lyn-dependent inhibitory tone by virus-like antigen display contributes substantially to this potency ^49^. Building on this, here we show that evasion of negative feedback downstream of Lyn also accounts for digital signaling and ultra-sensitive responses to pAg but not sAg, and results in divergent transcriptional programs (see model, **Figure S8D**).

Consistent with this framework, B cells exhibit analog signaling and robust affinity discrimination in response to sAg ^49^, properties that require negative feedback ^98–100^. In contrast, we previously showed that B cells lose affinity discrimination upon encounter with virus-like Ag display ^49^. We propose that this reflects a fundamental trade-off; digital signaling renders B cells insensitive to signal titration beyond a very low activation threshold; indeed, our imaging analyses suggest that one or two high valency virus-like particles may be sufficient to activate a B cell. Such sensitivity may have evolved as a first line of defense to mount a rapid immune response early in an infection, whereas affinity discrimination is re-imposed later in germinal centers, where inhibitory feedback is stronger ^101–103^.

One mechanism by which pAg evades Lyn-dependent signaling may be spatial re-organization of inhibitory co-receptors. Imaging studies have shown that B cell encounter with Ag on a membrane produces a micron-scale, synapse-like structure ^59^ within which CD19 is recruited ^104,105^, while inhibitory co-receptors such as FcγrIIb and CD22 are progressively excluded ^59,106^. Exclusion of the bulky transmembrane phosphatase CD45 from immune synapses may contribute to such membrane re-organization in B cells ^59,107,108^. However, immune synapses form on a longer time scale than the rapid, robust calcium responses we observe, suggesting additional mechanisms must operate. The size, epitope spacing, and conformational flexibility of pAg mirrors and may favor engagement of ∼100nm BCR nanoclusters on naïve B cells, while limiting access to inhibitory co-receptors ^109,110^.

pAg evades both Lyn and CD22-mediated inhibition. We find that CD22 partially mediates Lyn-dependent suppression of B cell responses to soluble stimuli, but its deletion does not fully phenocopy Lyn-deficiency. This may in part reflect reduced IgM BCR expression on MD4.CD22−/− cells, but we suspect additional Lyn substrates account for this discrepancy. Candidates include other ITIM co-receptors, the effector PTPases SHP-1, SHP-2, and the PIP_3_ lipid phosphatases SHIP-1 and PTEN. Consistent with this, we previously identified PIP_3_ as a key node of signal amplification downstream of pHEL ^49^ and deletion of lipid phosphatases is sufficient to activate anergic B cells ^26,27^. Mono-phosphorylation of ITAMs in the TCR and BCR can also recruit inhibitory phosphatases ^111,112^. It is alternatively possible that pAg engagement of BCR clusters could produce efficient dual ITAM phosphorylation, while sAg may result in mono-ITAM phosphorylation.

Distinct signaling by pHEL, sHEL, and anti-IgM may reflect distinct engagement of IgM and IgD BCR isotypes co-expressed on naïve B cells. IgM and IgD exhibit differential responses to antigen valency; rigid IgM hinge is argued to confer sensitivity to monovalent Ag, while flexible IgD hinge requires multivalent Ag for stimulation ^113^. However, this model remains controversial given evidence that IgD can respond to monovalent sHEL ^66,114^. IgD and IgM reside in distinct nanoclusters and may associate with distinct co-receptors ^109,110,115^, raising the possibility that antigen format selectively engages distinct receptor pools. Indeed, IgM but not IgD downregulation on anergic B cells is now well-recognized in both mice and humans ^115^. Selective restraint of IgM signaling by Lyn-dependent regulatory pathways may reinforce B cell tolerance, whereas preferential engagement of IgD by virus-like Ag could evade inhibitory tone and re-awaken anergic B cells.

We previously identified robust NF-κB activation by particulate but not soluble stimuli. Here we show that both Ag formats are linked to NF-κB through canonical Ca^2+^-and DAG-dependent activation of PKCβ ^68^. However, these two pathways are not equally limiting; although the Ca^2+^ threshold for NF-κB activation is high, addition of ionomycin failed to enhance NF-κB activation by soluble stimuli. In contrast, manipulation of DAG — using PMA or DGK inhibition — augments NF-κB activation by soluble stimuli, indicating that DAG rather than Ca^2+^ is limiting. This highlights DAG abundance as a crucial tipping point between tolerance and immunity in B cells. Consistent with this, transcriptional induction of *Dgk* expression in T cells is a core feature of anergy; indeed *Dgks* are a direct target of Egr2 ^116^. Their deletion is sufficient to overcome T cell tolerance, and these enzymes are an emerging target for cancer immunotherapy ^84,85,117^.

Unexpectedly, we find that nuclear NFAT is induced more strongly by soluble stimuli despite more robust Ca²⁺ responses to particulate antigen. A low threshold of cytosolic Ca^2+^ is sufficient for NFAT translocation, while high frequency oscillations with a long duty cycle maximize nuclear NFAT ^71,72,118,119^ – as in anergic B cells characterized by low amplitude Ca^2+^ oscillations ^32,35^. Indeed, superimposed addition of ionomycin did not further boost nuclear NFAT triggered by these stimuli. However, Ca^2+^ dynamics did not fully account for paradoxically reduced nuclear NFAT following pAg. Instead, we identify a DAG-dependent pathway that suppresses nuclear NFAT2 in B cells. Notably, engaging NF-κB downstream of CD40 does not suppress NFAT2, suggesting a role for PKC itself or PKC-dependent MAPK pathways as a possible link. JNK can antagonize NFAT nuclear translocation in T cells^120,121^, and may play this role in pAg-stimulated B cells.

It is well established that signal 1 (antigen) in the absence of signal 2 (CD28 co-stimulation) produces T cell anergy, a process governed by NFAT in the absence of AP-1 to drive transcriptional induction of a unique program of negative regulators in T cells ^30,65^. This includes E3 ubiquitin ligases (*Cblb*, *Itch*, and *Grail),* NFAT-dependent transcription factors (*Nr4as*, *Egr2* and *Egr3)* ^67^, and diacylglycerol kinase α (*Dgka*) which selectively suppresses MAPK and NF-κB signaling to re-enforce this program ^69,84,85^. This transcriptional program has many correlates in B cell anergy, including constitutive nuclear NFAT and impaired activation of JNK and NF-κB ^32,66^. B cell anergy is similarly characterized by upregulation of NFAT-dependent *Egr2/3*, deletion of which unleashes humoral immunity ^31^. We find that sHEL-stimulation *in vitro* is sufficient to recapitulate much of the transcriptional program seen in bona fide anergic B cells (MD4/ML5 model) ^66,87^. In contrast, pHEL largely evades this program. This is evident at the level of Egr2, the T cell-anergy marker FR4/*Izumo1r*, inhibitory co-receptor Tim3, and others. We propose that robust MAPK and NF-κB activation by pAg redirects NFAT away from tolerogenic targets and toward immunogenic gene programs.

It has been suggested that there is a limited time window for sAg-stimulated cells to obtain T cell help and escape apoptotic fate driven by accumulating Ca^2+^ ^25^. However, despite delivery of concurrent and maximal NF-κB activation, CD40 co-stimulation only partially suppressed the anergy-like transcriptional program induced by sHEL. We speculate that this may reflect the high and sustained nuclear NFAT levels driven by sHEL. By contrast, pHEL delivers a strong, stand-alone mitogenic stimulus to B cells, recapitulating the LPS-induced Myc program, yet does so through BCR-dependent PKC activation rather than the Myd88 pathway. The overlapping transcriptomes represent two distinct paths to a common immunogenic response (NF-κB with little NFAT), suggesting that in some ways virus-like antigen display is interpreted by the B cell as a danger signal despite being sensed through the BCR.

Functionally, sAg and pAg responses diverge in vitro and in vivo. Neither stimulus induces the robust survival programs associated with LPS or CD40, and both remain BAFF-dependent. However, sAg induces higher expression of pro-apoptotic transcripts such as Bcl2l11 (BIM), whereas pAg supports greater survival in vitro. Under physiological BAFF conditions in vivo, sHEL produces rapid B cell death, whereas pHEL immunization (with low Ag dose and neither adjuvant nor T cell help) results in expansion and survival at early time points, and productive Ab responses at later time points ^47,49^.

Particulate Ag induces much greater expression of CD86 than matched concentrations of soluble Ag, reflecting signaling potency, yet pMHC scales with Ag abundance, suggesting efficient delivery of signal 2. In contrast, anergic B cells exhibit impaired CD86 induction and therefore may promote T cell anergy ^122,123^. Indeed, delivery of physically linked peptide internal to pAg efficiently activates Ag-specific CD4 T cells, supporting not only T-independent (TI), but also T-dependent (TD) immune responses. It will be critical to determine how this liposomal platform regulates the balance between extra-follicular and germinal center responses since its modular design offers the capacity to independently program epitope density for BCR stimulation and cargo in the form of nucleic acids and/or peptides for vaccine development.

Collectively, our work explores the molecular basis for distinct B cell responses to antigen displayed in two different biophysical formats that mimic self-like or pathogen-like stimuli. These studies uncover how B cells are intrinsically wired to sense viral-like antigenic display as immunogenic, enabling mature B cells to remain inert to self-antigen yet respond robustly to self-mimicking pathogens.

## Materials and Methods

### Mice

IgHEL (MD4) mice were previously described ^50^. C57BL/6 and CD45.1+ BoyJ mice were originally from The Jackson Laboratory. MD4.*Lyn−/−* mice were generated by crossing MD4 mice to *Lyn−/−* mice, previously described ^124^. MD4. *CD22−/−* mice were generated by crossing MD4 mice to *CD22−/−* mice, previously described ^61^. sHEL Tg (ML5) mice ^50^ and OT-II TCR transgenic mice ^125^ were previously described. All strains were fully backcrossed to the C57BL/6J genetic background for at least 6 generations. Mice were used at 6-9 weeks of age for all functional and biochemical experiments. All mice were housed in a specific pathogen-free facility at University of California, San Francisco, according to the University Animal Care Committee and National Institutes of Health (NIH) guidelines. The study received ethical approval from the UCSF Institutional Animal Care & Use Program (Protocol number: AN201297).

### sHEL Reagents

Recombinant HEL-WT, HEL-D (R73E, D101R), HEL-T (R73E, D101R, R21Q) proteins were overexpressed in *E. coli* and purified to >95% purity following our established protocols as previously described ^48,49^ and are used to generate SVLS as described below. Recombinant sHEL was soluble comparator used to generate RNAseq data set described below. Elsewhere, unless otherwise noted, we used WT soluble HEL from Sigma (1μg/ml throughout or other doses as noted).

*<u>HELD fused with the Eα_52-68_ peptide</u>* was generated by appending the following amino acid sequence to the C-terminus of HELD through cloning, EEFAKFASFEAQGALANIAVDKANLDVMK. The flanking sequences of the peptide contain residues shown to be important for processing by antigen-presenting cells. This peptide was followed by GGGCHHHHHH, which encodes the single-engineered cysteine for liposomal conjugation and also the hexahistidine tag to facilitate the purification of this protein in native form from E. coli. The HELD-Eα protein was purified following our established protocols ^48^ but with important modifications. Specifically, after Ni-NTA column, the fractions containing HELD-Eα proteins were pooled, diluted with Buffer C (10 mM Na_2_HPO_4_, pH 7.0 at 22°C) by 10-fold in volume and loaded onto a HiTrap heparin column (GE) at a flow rate of 1.0 ml/min. The column was washed with Buffer C to baseline, and then sequentially washed with Buffer C containing 50, 100 and 150 mM NaCl respectively for 10, 30 and 45 min. The bound HEL proteins were then eluted in a gradient from 150 to 1000 mM increasing concentrations of NaCl in Buffer C over 25 column volumes at a flow rate of 1 ml/min. The fractions containing HEL proteins were pooled, diluted with Buffer C to 50 mM NaCl and loaded onto an Enrich S column (Bio-Rad) at a flow rate of 1.0 ml/min. The column was washed with Buffer C to baseline, and then sequentially washed with Buffer C containing 50 and 100 mM NaCl respectively for 5 min. The bound HEL proteins were then eluted in a gradient from 150 to 1000 mM increasing concentrations of NaCl in Buffer C over 25 column volumes at a flow rate of 1 ml/min. At this stage, the eluted HEL-Eα proteins were >95% pure. The protein was then filtered through 0.45 µm sterile syringe filter, concentration determined by absorbance at 280 nm, flash frozen in liquid N_2_ and stored in −80°C freezer. The fusion of the Eα peptide did not change the extinction coefficient of the protein at 280 nm under denaturing conditions and an extinction coefficient of 3.85×10^4^ M^-1^cm^-1^ was used for calculation of protein concentrations. The final yield of this protein was in general 10-fold lower than HELD, suggesting the presence of this peptide substantially lowered the solubility of the recombinant protein.

### SVLS synthesis and quantification

SVLS used throughout this study were prepared following protocols previously described by us^48,49^. In brief, SVLS were constructed using nonimmunogenic lipids, with phosphatidylcholine and cholesterol comprising ≥ 95% of all lipids. We prepared unilamellar liposomes using a mixture of DSPC, DSPE-PEG maleimide, and cholesterol. Empty liposomes were generated using PBS and extruded membrane with a pore size of 100 nm. Hen egg lysozyme (HEL) recombinant proteins or affinity variants with free engineered cysteines were conjugated to the surface of liposomes via maleimide-thiol chemistry at specific engineered cysteine and programmable epitope density by modulating molar percentage admixed DSPE-PEG maleimide (0.5-5%). A large excess of free cysteines at the end of a 1 h cross-linking reaction was used to quench all of the available maleimide groups that remain on the liposomal surface. After this conjugation, the liposomes were purified away from free excess HEL proteins by running through a size exclusion column (SEC). Epitope density was quantified using methods that we established previously^48,126^ that were also validated by single molecule fluorescence ^127,128^. Control SVLS (pLIP02) had no surface protein conjugation and were used as controls in generation of RNAseq dataset discussed below.

*<u>Synthesis of fluorescent pHELD</u>* (pHELD-AF594, ED=127) harboring Alexa-594 dye was previously described and validated^49^ and were used for live cell microscopy described below.

*<u>Synthesis of SVLS with HELD-Eα_52-68_ peptide</u>* (pHELD-Eα, ED=99) as described above, except recombinant HELD with Ea peptide fusion was used for conjugation.

*<u>Synthesis of SVLS harboring free internal OVA_323-339_ peptide</u>* (pHEL-OVA, ED=9) was conducted by extrusion of the lipids in the presence of 1 mM OVA peptide dissolved in PBS followed by steps described above. The OVA peptide was custom synthesized by Biomatik with the following sequence, ISQAVHAAHAEINEAGR.

### Antibodies and Reagents

#### Antibodies for surface markers

Live/Dead NIR (Invitrogen), B220 (clone: RA3-6B2, BD or Tonbo) CD21 (clone: 7G6, BD), CD23 (clone: B3B4, eBioscience), IgM (clone: MA-69, BioLegend), IgM Fab’1 (polyclonal, Jackson ImmunoResearch), IgD (clone: 11-26, eBioscience), CD69 (clone: H1.2F3, Tonbo) Fr4 (clone: 12A5, BioLegend), Lag-3 (clone: C9B7W, eBioscience), Tim-3 (clone: RMT3-23, BioLegend), PD-1 (clone: 29F.1A12), Y-Ae (clone: eBioY-Ae (YAe, Y-Ae), eBioscience) CD86 (clone: GL-1, BioLegend), MHC II (clone: M5/114.15.2, Tonbo), Vb5.1(clone: MR9-4, BioLegend), Va2 (clone: B20.1, BioLegend). All commercial surface Abs were used at 1:200 dilution.

#### Antibodies for intracellular staining

Egr2 (clone: erong2, ThermoFisher), c-Fos (clone: T.142.5, Invitrogen), T-bet (clone: 4B10, BioLegend), pErk1/2 T202/Y204 (clone: 194G2, Cell Signaling), c-Myc (clone: D84C12, Cell Signaling). All intracellular antibodies were used at 1:100 or 1:200 dilution.

#### Antibodies for nuclei staining

NFATc1/NFAT2-PE (clone: 7A6, BioLegend), NFATc2/NFAT1-AF488 (clone: D43B1, Cell Signaling), NF-κB p65 mAb (clone D14E12, Cell Signaling) unconjugated, c-Rel-PE (clone: 1RELAH5, eBioscience). All nuclear staining antibodies were used at 1:100 dilution overnight.

#### Secondary antibodies for intracellular and nuclear staining

AffiniPure F(ab’)₂ Fragment Donkey Anti-Rabbit IgG (H+L) secondary Ab conjugated to APC was from Jackson Immunoresearch.

#### Stimulatory antibodies and reagents

Goat anti-mouse IgM F(ab’)2 (Jackson Immunoresearch), anti-CD40 (1μg/ml, hm40- 3 clone; BD Pharmingen), recombinant murine BAFF (20ng/ml, cat#2106-BF, R&D), LPS (10μg/ml O26:B6; Sigma), CpG (2.5μM ODN 1826; InvivoGen), phorbol myristate acetate (PMA; 0.2-20ng/mL, Sigma), Ionomycin (0.01-1μM, Calbiochem).

### Media

Complete culture media was prepared with RPMI-1640 + L-glutamine (Corning-Gibco), Penicillin Streptomycin L-glutamine (Life Technologies), 10 mM HEPES buffer (Life Technologies), 55 mM β-mercaptoethanol (Gibco), 1 mM sodium pyruvate (Life Technologies), Non-essential Amino acids (Life Technologies), and 10% heat inactivated FBS (Omega Scientific). This was used for all *in vitro* culture and stimulation except for calcium signaling experiments. Calcium flux media was prepared with RPMI, HEPES, PSG as above and 5% FCS.

### Inhibitors

Ly294002 (PI3Ki): 1-100mM as noted in figure panels (Calbiochem), Bay61-3606 (Syki): 0.3-2.5mM as noted in figure panels (Calbiochem), Ibrutinib (Btki): 3 −100 pM as noted in figure panels (gift from Jack Taunton, UCSF), PP2 (SFKi): 10mM (Calbiochem), EGTA 2mM (Sigma), Cyclosporin A (CSA): 1μM (Calbiochem), U0126 (MEKi): 100μM (MedchemExpress), R59949 (DGKi): 100μM (MedchemExpress), Go6976 (PKCi): 10μM (MedchemExpress), R568 (IRAK1/4i): 1µM (gift from Rigel).

### Cellular staining for flow cytometry

Cells were stained with LIVE/DEAD Fixable NIR (Invitrogen) and surface marker antibodies in PBS for 30 minutes on ice at 1×10^6^ cells/well in a 96-well plate. Cells were washed with FACS Buffer [PBS (Gibco) + Penicillin Streptomycin L-glutamine (Life Technologies) + EGTA + 5% FBS] and resuspended in FACS buffer for analysis.

### Intracellular staining for flow cytometry

After surface marker and LIVE/DEAD staining (see above), cells were fixed with FoxP3 Fixation-Permeabilization Buffer (Invitrogen) for 20 minutes on ice, washed with Permeabilization Wash Buffer (Invitrogen), and stained with intracellular antibodies made up in Permeabilization Wash Buffer for 60 minutes RT or overnight at 4°C. Cells were then washed in Permeabilization Wash Buffer, and resuspended in FACS buffer for analysis.

### Intracellular staining to detect p-Erk and c-Myc

Following *in vitro* stimulation, cells were immediately fixed in 2% paraformaldehyde (PFA) for 15 minutes at RT, washed in FACS buffer, and permeabilized with ice-cold 90% methanol at −20°C overnight, or for at least 30 minutes on ice. Cells were then washed in FACS buffer, stained with intra-cellular antibody for 40 minutes on ice, washed in FACS buffer, and then stained with APC-conjugated secondary Goat anti-Rabbit antibody along with directly conjugated antibodies to detect surface lineage and/or subset markers for 40 minutes on ice. Samples were washed and then resuspended in FACS buffer for analysis.

### Vital dye loading

Cells were loaded with CellTrace Violet or CellTrace Yellow (CTV or CTY; Invitrogen) per the manufacturer’s instructions except at 5×10^6^ cells/mL rather than 1×10^6^ cells/mL.

### Flow cytometry data analysis, software, and statistics

After staining, cells were analyzed on a Fortessa (Becton Dickinson) or Aurora (Cytek). Data analysis was performed using FlowJo (v10.8.2) software (Treestar Inc.). Proliferative indices ‘division index’ and ‘% divided’ were calculated using FlowJo. Statistical analysis and graphs were generated using Prism v10 (GraphPad Software, Inc). Statistical tests used throughout are listed at the end of each figure. Student’s paired T test (two-tailed, parametric assuming gaussian distribution) was used to calculate p values for all comparisons of two pre-specified groups. One-way ANOVA was performed when more than two groups were compared to one another. Mean is plotted and error bars in graphs represent SEM. All statistical tests were two-sided.

### Intracellular calcium flux by flow cytometry

Cells were loaded with 5μg/mL Indo-1 AM (Life Technologies) per manufacturer’s instructions, stained with lineage markers in Calcium Flux Media for 15 minutes on ice and then resuspended at 10×10^6^ cells/mL. Cells were then rested at 37 C for 5 min, and Indo-1 fluorescence was measured by FACS immediately prior to and after stimulation to determine intracellular calcium. Analysis was performed in FlowJo by deriving a parameter of Indo-1 unbound/Indo-1 bound and generating kinetic plots.

### Live cell imaging of intracellular calcium entry by microscopy

A 96-well slide glass MatriPlate (cat#: BRKS-MGB096-1-2-LG-L, NETA Scientific) was coated overnight with 100uL/well of PBS with ICAM-1 (5μg/mL, SinoBiological) and fibronectin (10μg/mL, Millipore Sigma). B cells were isolated (see above) and loaded with Fluo-4 AM (Invitrogen), according to the manufacturer’s instructions. Cells were then stained with B220-AF647 on ice for 15 minutes in calcium flux media (see above). After washing in calcium flux media, B cells were plated at 500,000 cells/well on pre-coated slide glass plate, pre-warmed at 37C for 30 minutes, and placed on the Nikon A1R TIRF microscope, using climate control set at 37C and 5% CO_2_. Live calcium flux was acquired by adding 10μL of 10x stimulations to the well and acquiring images using a non-TIRF angle every 2 seconds for 5 minutes.

### Live cell calcium imaging analysis

B cells were identified as regions of interest (ROI) in Fiji using B220-AF647 staining. Fluo4-AM MFI was tracked over time in 2-second intervals for individual cells using TrackMate (version 8). Individual cell tracks were filtered for those with ≥ 30 records (snapshots of Fluo4AM fluorescence data covering at least 60 seconds). Single-cell metrics of calcium signaling for **Figure 2** were defined as follows: time to calcium entry is the earliest time record at which Fluo4AM MFI reached ≥ 1.5X the initial Fluo4AM MFI in a given cell, peak MFI is the maximum post-stimulation Fluo4AM MFI per cell, and area under curve is the sum of Fluo4AM MFI multiplied by 2 (2-second intervals between time records).

### Population kinetics analysis of B cell activation by pAg particles

#### Calcium traces processing for Figure S2

For each recording (defined by pAg concentration and replicate), we first filtered out spurious calcium fluorescence traces, excluding traces lasting fewer than 50 frames or beginning more than 30 s after the start of the recording. We retained only replicates containing at least 15 cells, to ensure meaningful fits could be performed. To distinguish B cells exhibiting cytosolic calcium increase above background, we defined a calcium fluorescence intensity threshold. We set the threshold as a factor (*f*) times the background fluorescence; the background was itself determined by finding the minimum fluorescence in each trace, then taking the median over traces (**Figure S2C**). We used the same factor *f* across recordings. We tested a range of *f* values between 1.1 and 2.5 and settled on a default value *f* = 1.5 (**Figure S2C**). We determined the activation time *t*^∗^ of each B cell by linear interpolation of the threshold crossing time. Traces already above threshold at the start of imaging were assigned *t*^∗^ = 0. Cells that never crossed the threshold were labeled inactive for the entire recording. Moreover, we normalized the activated cell distributions to the fraction of tracks that activate in the highest concentration (10 pM) replicates, reasoning that non-responders in this data set represent the proportion of dead/unresponsive cells.

#### Mathematical modeling of the distribution of B cell activation times (eCDF plots)

We treated B cell activation as a stochastic process governed by the waiting time for a B cell to encounter one or more randomly diffusing particles and subsequently reach the calcium activation threshold. The activation time *t*^∗^ thus corresponds to the first-passage time (FPT) for the B cell to reach the “active” state. The cumulative distribution function (CDF) of the FPT, 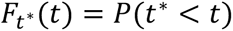, gives the probability that activation has occurred at or before time *t*. The CDFs from different concentrations of pHEL were shown in **Figure 2H** and **Figure S2D**.

#### Kinetic analysis to determine effective activation rate

We first sought to determine how particle concentration affected the overall rate of B cell activation. pHEL CDFs of empirical activation times were first fit to a single-exponential function CDF 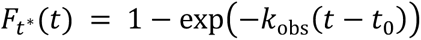 using maximum likelihood, as described in more detail below (**Figure S2D**). The parameter *t_0_* is fitted along with *k*_obs_ to account for uncertainty in the time origin of the experiment. The resulting optimal values of *k*_obs_ were plotted as a function of pHEL concentration (**Figure S2E**) and compared to the diffusion-limited particle binding rate expected from physical principles, *k_diff_* = κ*c*, with

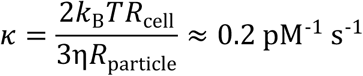

where η is the viscosity of the complete media which we approximate using the value of water, 0.7 mPa of water at *T* = 37^∘C^ = 310 K, *R*_cell_ = 5 µm is a typical B cell radius, and *R*_particle_ = 60 nm is the typical radius of virus-like particles, including pHEL generated for our studies^47^. We anticipated that activation at low concentrations would be limited by particle diffusion and have *k*_obs_ close to κ*c*. Indeed, we found maximum likelihood estimates *k*_obs_ = 0.0041 s^-1^ and 0.0054 s^-1^ for the two 0.01 pM replicates, comparable to theoretical κ*c* = 0.002 s^-1^ calculated above (despite viscosity difference of media), and also to experimentally measured rate for binding of single HIV-1 virions ^64^, indicating that particle diffusion rate is limiting for B cell activation at very low particle concentrations.

#### One-hit and multi-hit models of B cell activation

We developed *M*-hit models of B cell activation incorporating 1, 2, or 3 particle binding steps with hit rate *k_1_* followed by an intrinsic cell activation step with rate *k_2_*, which can respectively become limiting in low and high particle concentration conditions (**Figure S2F**). We assumed negligible off-rate given pHEL with epitope density > 100. We derived analytical expressions for the FPT CDF and probability density function (PDF) of each model using Laplace transform methods. For instance, the one-hit model has

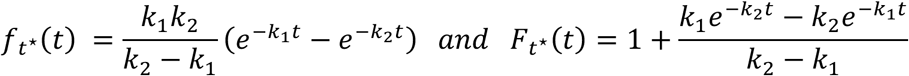

with similar, albeit longer, expressions for the 2-hit and 3-hit models. These models predict distinct CDF curves at low particle concentrations; we could therefore fit them to low concentration datasets to determine whether one or more particle contacts are required to explain B cell activation. For this purpose, we again introduced a time origin parameter *t*_0_, implemented by replacing *t* with *t* − *t*_0_ throughout the PDF and CDF expressions.

#### Maximum likelihood parameter estimation

We tested these models by fitting to the empiric activation times CDF, treating individual B cells as independent realizations of the same first-passage process, and inferred parameters via maximum likelihood. To extract the observed activation rate *k*_obs_, we fitted *k*_obs_ and *t_0_* on each dataset separately. For the 1-hit, 2-hit, and 3-hit models, we inferred one global cell-intrinsic activation rate *k_2_* across all datasets, and one pair (*k_1_*, *t_0_*) per dataset, to account for varying concentrations and replicate variability. We therefore maximized a total log-likelihoodlog ℒ_tot_ = ∑_j_ log ℒ_j_ across datasets. For each dataset, we used a likelihood function ℒ_;_ including a PDF term for each activation event up to a cutoff time *t*_cut_ – chosen as the time at which the empiric CDF reaches 75% active cells – and a CDF term for later events. This cutoff reduced sensitivity to spurious inactive tracks not corrected by our eCDF normalization while including most activation events in PDF terms. To define this likelihood, consider that recording *j* contains *N* cells with valid calcium traces, and suppose *K*_0_ tracks are already activated at *t=0*, *K_1_* cells become activated at times 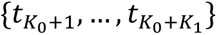 between *t=0* and *t*_cut_, and the remaining _N_ – *K*_0_ – *K*_1_ cells are still resting at *t*_cut_. Initially active cells contribute 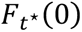 to the likelihood, and cells still resting at *t*_cut_ contribute 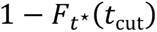. The log-likelihood for dataset *j* is therefore

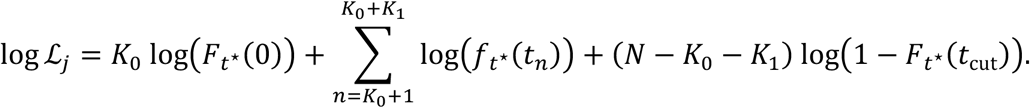

We maximized the log-likelihood with respect to parameters using the interior-point Newton algorithm in the *Optim.jl* module in Julia ^129,130^. For **Figure S2G-I**, we repeated this process across a range of fluorescence threshold factors *f*. The fits for *k_1_* were generally insensitive to *f*, whereas *t_0_* varied more because the earliest activation time *t_1_* is delayed as *f* increases. Our code for this analysis was deposited on Zenodo at https://doi.org/10.5281/zenodo.19657273.

### In vitro B cell culture and stimulation

Splenocytes or lymphocytes (as noted in legends) were harvested into single cell suspension through a 40μm cell strainer, subjected to red cell lysis using ACK (ammonium chloride potassium) buffer in the case of splenocytes, +/– CTV loading as described above, and plated at a concentration of 1-10 × 10^5^ cells/200 μL (depending on assay) in round bottom 96 well plates in complete RPMI media with stimuli for 0-4 days. In vitro cultured cells were stained to exclude dead cells as above in addition to surface or intracellular markers for analysis by flow cytometry depending on assay.

### B and CD4+ T cell benchtop purification

Purification was performed utilizing MACS separation per manufacturer’s instructions. In short, pooled spleen and/or lymph nodes were prepared utilizing the B cell or CD4+ T cell kits (Miltenyi) and purified by negative selection through an LS column (Miltenyi).

### Nuclei isolation

The nuclei isolation protocol was adapted from online sources (National Cancer Institute Experimental Transplantation and Immunology Branch Flow Cytometry Core Laboratory; home.ccr.cancer.gov/med/flowcore) and previously published reports ^131^. In brief, purified splenic/LN B cells (protocol described above) were harvested from mice and pre-warmed in a 96-well plate for 10 min at 37°C with 1.5×10^6^cells/100 μL per well in complete media. Cells were stimulated with 100 μL of 2x stimuli for 20 min at 37°C. Stimulated cells were spun at 300 x *g* at 4°C, and the pellets were immediately resuspended with 200 μL of ice-cold Buffer A containing 320 mM sucrose, 10 mM HEPES (Life Technologies), 8 mM MgCl2, 1× EDTA-free complete Protease Inhibitor (Roche), and 0.1% (v/v) Triton X-100 (Sigma-Aldrich). After 15 min on ice, the plate was spun at 2000 × *g* and 4°C for 5 min. This was followed by two 200 μL washes with Buffer B (Buffer A without Triton X-100) and spinning at 2000 × *g* and 4°C for 5 min. After the final wash, pellets were resuspended with 200 μL Buffer B containing 4% paraformaldehyde (electron microscopy grade; Electron Microscopy Sciences), and nuclei were rested on ice for 30 min for fixation. The nuclei were spun at 2000 x *g* and 4°C for 5 min, followed by two washes at 1000 × *g* and 4°C for 5 min and resuspension in stain.

### Nuclei staining

Isolated nuclei were washed with 200 μL nuclei Perm Buffer (nuclei FACS Buffer with 0.3% Triton-X 100) and spun at 1000 × *g* and 4C for 5 min. Isolated nuclei were stained with 50 μL intracellular antibody cocktails (see above) overnight at 4C. Nuclei were spun at 1000 x *g* and 4°C for 5 min, washed withi FACS Buffer and stained with 50 μL anti-rabbit IgG-APC (see above) diluted in Perm Buffer for 60 min on ice in the dark. The nuclei were spun at 1000 x *g* and 4°C for 5 min, washed with nuclei FACS Buffer, and resuspended in FACS Buffer and analyzed immediately on a BD Fortessa or Cytek Aurora cytometer.

### RNAseq sample preparation

B cells were isolated from MD4 lymph nodes via bench-top negative selection (as described above) and stimulated overnight (16 hours) at 37C at 1×10^6^ cells/well in a sterile 96-well plate. Biological triplicate samples were stimulated with recombinant sHEL (1μg/mL), recombinant sHEL with anti-CD40 Ab (100ng/mL), pHELT (ED=243) or LPS (10μg/mL). Unstimulated samples were harvested at 2 hours to circumvent poor viability at later timepoints. A small aliquot was taken for flow cytometry staining to validate activation and the remaining culture was transferred to sterile, RNAse-free Eppendorf tubes. 1mL of cold PBS was added and cells were spun at 4C. Supernatant was removed and pellets were snap frozen using a liquid nitrogen bath, stored at −80 C, and then sent for further processing and sequencing at Novogene. In brief, mRNA was purified from total RNA using poly-T oligo-attached magnetic beads. After fragmentation, the first strand cDNA was synthesized using random hexamer primers followed by the second strand cDNA synthesis. After library prep, sequencing was performed with NovaSeq X Plus Series (PE150).

### RNAseq data processing and analysis

Raw fastq files were adapter- and quality-trimmed with Trim Galore v0.6.10 and then mapped to the GRCm39 (mm39) genome with the GENCODE vM36 gene annotation using STAR v2.7.11b ^132^. PCR duplicate removal and gene-level counts generation were handled by STAR with the options --bamRemoveDuplicatesType UniqueIdentical and --quantMode GeneCounts, respectively. Counts were normalized to transcripts per kilobase per million (TPM). Differentially expressed genes between sample groups within the data set were determined using DESeq2 ^133^, v1.50.0, and genes that exhibited absolute log2(fold change) ≥ 1 and Benjamini–Hochberg false discovery rate (FDR)-corrected *p*-value ≤ 0.05 were considered significant. All plots were generated using Seaborn v0.13.2 or matplotlib v3.10.7. Data is available in **Supplementary Table 1a**.

### Principal Component Analysis

Principal component analysis (PCA) was performed on variance-stabilized gene-level RNAseq counts, using the experimental design to calculate dispersion, with DESeq2 v1.50.0 (Fig 5D, E) and PCAtools v2.18.0 R (Fig S5B) packages. PCA loadings were derived for the top 5 principal components in the PCA excluding unstimulated samples, and the top ≤ 16 loadings (genes with the most positive or most negative contributions to a specific component) were plotted in Figure S5B. Heatmaps were generated using the Morpheus online tool (https://software.broadinstitute.org/morpheus/) with default clustering of rows (Pearson minus one) unless otherwise noted in figure legend. PCA loadings available in **Supplementary Table 1b**.

### Gene set enrichment analyses

Pairwise gene set enrichment analyses were performed with GSEA v4.3.3^134,135^ on gene-level TPM data. Included in the analysis were MSigDB Hallmark gene sets and B cell-specific gene sets filtered for absolute log2(fold change) ≥ 1 and *p*-value or FDR ≤ 0.05; split into significantly upregulated or downregulated gene sets when applicable. GSEA datasets and results available in **Supplementary Table 1c, 1d**.

### In vitro B cell-T cell co-culture

CD4+ T cells were isolated from the spleen and lymph nodes of OT-II mice utilizing MACS separation and the CD4+ T cell negative selection kit (Miltenyi, see above). Isolated OT-II T cells were then CTV labeled (see above) plated in a 96-well round bottom plate at a ratio of 1:1 with isolated MD4 B cells or polyclonal APCs for a total of 1×10^5^ cells/well. Stimuli (pHEL10pM, pHEL-OVA_323-339_ 10pM, OVA_323-339_ 5μM [Genscript]) were added +/− rmIL-2 (5ng/mL, Peprotech) and cultures were placed at 37C for 1-4 days to assess T cell activation and proliferation.

### Adoptive transfer of MD4 to sHEL hosts

MD4 splenocytes were isolated and loaded with Cell Trace Violet vital dye and 5×10^6^ cells injected into either WT (C57BL/6) or sHEL-Tg (ML5) recipient hosts via tail vein. Two days post injection, splenocytes were harvested from host mice and analyzed by flow cytometry.

### Adoptive transfer of MD4 to WT hosts + pHEL immunization

Lymphocytes from MD4 mice were harvested, loaded with Cell Trace Violet vital dye as noted, and 5×10^6^ cells were transferred via tail vein injection into CD45.1+ C57BL/6 hosts. The following day, 200mL of 0.125nM pHELT (ED 243) in PBS (∼0.1mg HEL protein) was administered with *<u>no</u>* adjuvant via tail vein injection. Unimmunized hosts with transferred cells served as controls. Three days later, host splenocytes were harvested for analysis via flow cytometry.

## Supporting information

Video S1 ionomycin

Video S2 pHEL-AF594 10pM

Video S3 pHEL-AF594 1pM

Video S4 pHEL-AF594 0.1pM

Video S5 pHEL-AF594 0.01pM

Video S6 sHEL 1ug/mL

Video S7 sHEL 0.1ug/mL

Video S8 sHEL 0.01ug/mL

Video S9 sHELD 1ug/mL

Video S10 sHELD 0.1ug/mL

Video S11 sHELD 0.01ug/mL

Video S12 anti-IgM 10ug/mL

Video S13 anti-IgM 1ug/mL

Video S13 anti-IgM 0.1ug/mL

Video S15 pHEL-AF594 1pM zoom

Video S16 pHEL-AF594 1pM zoom 2

Video S17 pHEL-AF594 1pM zoom 2

## Data availability

Raw Fastq and processed data files (either RPKM or TPM) for the original RNA-seq analyses have been deposited in the NCBI Gene Expression Omnibus and are publicly available under accession code GSE285400 and DEG are also provided in **Supplementary Table 1a**. All data are included in the Supplementary Information or available from the authors, as are any unique reagents used in this article

## Acknowledgments

We thank our funders – NIAID R01AI155653 (WC, JZ), R01AI148487 (JZ), R01AI165706 (JZ), R01AI114575 (JZ), NIH Training Grant T32 AI007334 (JR, JL), UCSF IRACDA (AR). ARM was supported in part by an NIH/NIGMS T32 GM145304 Cellular Biotechnology Training Program, a predoctoral fellowship from the American Foundation for Pharmaceutical Education, and a University of Michigan Rackham Predoctoral Fellowship. NWW was supported in part by Princeton University through the Center for the Physics of Biological Function. We thank Al Roque for help with mouse husbandry. We thank Dr. Roger Sciammas for his kind suggestion regarding Eα peptide sequence for fusion with HELD. We thank Dr. Jeremy F. Brooks and Dr. Wan-Lin Lo for critical reading of the manuscript.

