## Supplementary material for "Virus-like antigen display delivers a stand–alone danger signal through the BCR that circumvents tolerance": Video Legends

**Videos S1-17. B Cell Ca2+ responses to stimuli.** Live microscopy imaging of purified B cells (B220-AF647+, red) loaded with Fluo4-AM Ca^2+^ indicator dye (green) responding to various stimuli as noted below, including pHEL-AF594 (pink). S15-17 depict close-up examples of single B cells stimulated with 1pM pHEL that show particle encounter and subsequent B cell response. All images captured at 100x. All videos represent images/frames captured every 2 seconds for 5 minutes; Videos are compressed to 7 frames/second (300 sec 🡪 21 sec which corresponds to 14x time compression).

**Video S1: B cell Ca^2+^ responses to** 1μM **ionomycin**

**Video S2: B cell Ca^2+^ responses to pHEL-AF594 10pM**

**Video S3: B cell Ca^2+^ responses to pHEL-AF594 1pM**

**Video S4: B cell Ca^2+^ responses to pHEL-AF594 0.1pM**

**Video S5: B cell Ca^2+^ responses to pHEL-AF594 0.01pM**

**Video S6: B cell Ca^2+^ responses to sHEL 1μg/mL**

**Video S7: B cell Ca^2+^ responses to sHEL 0.1μg/mL**

**Video S8: B cell Ca^2+^ responses to sHEL 0.01μg/mL**

**Video S9: B cell Ca^2+^ responses to sHELD 1μg/mL**

**Video S10: B cell Ca^2+^ responses to sHELD 0.1μg/mL**

**Video S11: B cell Ca^2+^ responses to sHELD 0.01μg/mL**

**Video S12: B cell Ca^2+^ responses to anti-IgM 10μg/mL**

**Video S13: B cell Ca^2+^ responses to anti-IgM 1μg/mL**

**Video S14: B cell Ca^2+^ responses to anti-IgM 0.1μg/mL**

**Video S15: Single B cell Ca^2+^ responding to pHEL-AF594 1pM**

**Video S16: Single B cell Ca^2+^ responding to pHEL-AF594 1pM**

**Video S17: Single B cell Ca^2+^ responding to pHEL-AF594 1pM**
