## Supplementary Legends and Figures 1 to 8 for "Virus-like antigen display delivers a stand–alone danger signal through the BCR that circumvents tolerance"

#### Supplementary Figure Legends

**Supplementary Figure 1:**

**S1A:** Crystal structure of sHEL protein (1C08 pdb) highlighting point mutations that reduce affinity for the Hy10 BCR.

**S1B:** Model depicting kinase inhibitor targets that mediate proximal BCR signal transduction.

**S1C:** Intracellular pErk induction measured by flow cytometry 20 minutes post stimulation. MD4 pooled splenocytes and lymph node cells were pre-incubated with dose titrations of Syk inhibitor (Bay61-3606, 2.5, 1.25, 0.6, 0.3μM) or Btk inhibitor (Ibrutinib, 100, 30, 10, 3pM) followed by stimulation with sHEL (1μg/mL) or pHEL (1pM). Histograms depict B220+ cells and are representative of at least three independent experiments.

**S1D:** $IC_{50}$ values calculated from inhibitor dose titration experiments as in **1D, 1E** and **S1C**. Each linked dataset represents an individual experiment. $IC_{50}$ was calculated by plotting dose response data and performing a non-linear regression analysis. Each data point represents a biological replicate. P values acquired from Student's paired t-test.

**S1E:** MD4 pooled splenocytes and lymph node cells were loaded with Indo-1 $Ca^{2+}$ indicator dye and stained for viability and surface markers. Cells were run in real time by flow cytometry to assess intracellular $Ca^{2+}$ entry after stimulation with ionomycin 1μM, anti-IgM 10μg/mL, sHEL 1μg/mL or pHEL 1pM with or without addition pan Src family kinase (SFK) inhibitor PP2 (10μM). Data representative of three independent experiments.

**S1F:** MD4, MD4.*Lyn-/-* and MD4.*CD22-/-* pooled splenocytes and lymph node cells were stained for viability, B220, CD23 and CD21. Top: percentage of B220+ cells of live cells. Bottom: percentage of live B220+ that are CD23+ $CD21^{low}$ (follicular B cells), $CD23^{low}$ $CD21^{high}$ (marginal zone B cells) or CD23- CD21- (transitional B cells).

**S1G:** MD4, MD4.*Lyn-/-* and MD4.*CD22-/-* pooled splenocytes and lymph node cells were stained for viability, B220, IgD and IgM. Histograms depict MFI of IgM and IgD on live B220+ cells.

**S1H:** Histogram showing MFI of IgM-Fab'1 of various B cell subsets from MD4 and MD4.*CD22-/-* live B220+ splenic and lymph node cells.

**S1I, J:** Intracellular $Ca^{2+}$ entry in MD4 and MD4.*CD22-/-* $CD23^{hi}$ live B220+ cells was assessed by flow cytometry after stimulation with ionomycin 1μM, anti-IgM 10μg/mL, sHEL 1μg/mL or pHEL 1pM. CD23+ B220+ cells from MD4.*CD22-/-* mice are subgated by IgM-Fab'1 surface level expression (as in **1I**). Data representative of two independent experiments.

**S1K, L:** Erk phosphorylation in MD4 and MD4.*CD22-/-* $CD23^{hi}$ B220+ cells was assessed by intracellular staining after stimulation with PMA 20ng/mL, anti-IgM 10μg/mL, sHEL 1μg/mL or pHEL 1pM for 20 minutes. CD23+ B220+ cells from MD4.*CD22-/-* mice are subgated by IgM-Fab'1 surface level expression, stained post fixation and permeabilization (as in **1K**). Data representative of two independent experiments.

**Supplementary Figure 2**. **Calcium microscopy and population kinetics analysis of B cell activation by VLPs.**

**S2A:** Snapshots from calcium video microscopy recordings for two different VLP concentration conditions.

**S2B:** Analysis pipeline for live $Ca^{2+}$ microscopy. B cells were identified as regions of interest (ROI) in Fiji using B220-AF647 staining. TrackMate was then used to track ROIs over time, tracking B cells even as they floated across the field of view. Mean fluorescence intensity of Fluo4-AM for each ROI at each timepoint was recorded and exported as a csv file for further analysis.

**S2C:** Example of calcium fluorescence intensity traces (1pM pHEL-AF594) and thresholding to determine B cell activation times. The threshold is set to be a factor *f* times the background fluorescence in each recording (1.8 by default).

**S2D:** Particle CDFs were fit with an exponential distribution function $1 - \exp(-k_{\mathrm{obs}}(t - t_0))$ by maximum likelihood (Methods), to extract effective activation rates. One replicate per concentration is shown for clarity.

**S2E:** Plots of resulting $k_{obs}$ values obtained from fitting in **S2D** versus particle concentration. The dashed line represents the theoretical diffusion-limited particle binding rate; $k_{diff} = \kappa c$, with $\kappa = \frac{2k_{\rm B}TR_{\rm cell}}{3\eta R_{\rm particle}} \approx 0.2\ {\rm pM}^{-1}\ s^{-1}$ (see methods). The results suggest that particle binding limits activation at low concentrations, while a cell-intrinsic step limits activation at higher concentrations (inset schematic).

**S2F:** Cell activation models where one, two, or three particle binding steps (``hits”) with rate $k_1$ are needed before a cell can undergo a final activation step with rate $k_2$ (insets; we neglect reverse reactions as described in methods).

**S2G:** Example model fits on a low-concentration replicate (0.01 pM pHEL), where the three models have distinct curve shapes and root-mean-squared errors (RMSE). Also, a time origin parameter $t_0$ captures the uncertainty in the precise recording start time and underlies a potential lag time before activation kinetics start. Each model is fitted by maximum likelihood, with a global rate $k_2$ across datasets and one value of $k_1$ and $t_0$ per replicate (Methods).

**S2H**: As in **S2G**, for a high-concentration replicate (1 pM pHEL), where all models have similar kinetics limited by the cell-intrinsic step with rate $k_2$.

**S2I:** Maximum likelihood estimates of the binding rate $k_1$ for the 1-, 2-, and 3-hit models for each low-concentration replicate (0.01pM a, b and 0.1pM a, b), as a function of the choice of fluorescence threshold factor *f*. Most fits of *k* do not depend strongly on *f* in a range between 1.25 and 2.2 (grey area).

**S2J:** Comparison of the fit quality of the 1- or 2-hit models, across datasets (dot colors) and choice of threshold factor *f* (dot sizes).

**S2K**: As in **S2J**, comparing 3-hit model to 1- or 2-hit models.

**Supplementary Figure 3:**

**S3A:** Immgen gene skyline data demonstrating relative abundance of *NFATc2* (NFAT1, right) and *NFATc1* (NFAT2) transcript in B and T lymphocytes.

**S3B:** Nuclear staining of NFAT1 and NFAT2 in B cells following 20-minute stimulation with denoted stimuli.

**S3C:** Nuclear staining of NFAT1, NFAT2 and p65 in CD4+ T cells and B cells after 20 minutes stimulation with noted stimuli. Dotted and solid lines denote baseline and maximal, respectively, gMFI for CD4+ T cells for each transcription factor.

**S3D:** Quantification of raw gMFI data from **S3C**. P values acquired via student's paired t test.

**S3E:** NFAT2 staining in B cell nuclei after stimulation with and without 30-minute preincubation with 1μM EGTA. Histograms representative of three experimental replicates.

**S3F:** NFAT2 in B cell nuclei after stimulation with and without 30-minute preincubation with Cyclosporin A at 1μM. Histograms representative of two experimental replicates.

**S3G-H:** Quantification of data in **S3E** and **S3F**. Each linked set of datapoints represents an experimental replicate. P values were acquired by performing paired Student's t-test.

**S3I:** Intracellular c-Fos staining in whole B cells after 2 hours stimulation with and without 30-minute preincubation with MEK inhibitor U0126 at 100μM. Histograms representative of two independent experiments.

**S3J-K:** Ratio of mean normalized c-Fos gMFI to mean normalized nuclear NFAT2 gMFI **(S3J)** or pErk gMFI **(S3K)** across various stimulation conditions.

**Supplementary Figure 4**

**S4A:** Schematic depicts small molecule inhibitors deployed to selectively perturb DAG and calcium-dependent pathways in B cells.

**S4B:** p65 staining in B cell nuclei. B cells were incubated with PKC inhibitor (Go6976 10μM) for 30 minutes prior to stimulation for 20 minutes. Data representative of at least four independent experiments.

**S4C:** p65 staining in B cell nuclei. B cells were incubated with EGTA (2 mM) for 10 minutes prior to stimulation for 20 minutes. Data representative of at least four independent experiments.

**S4D-E:** Quantification of data in **S4B** and **S4C.** gMFI values were normalized to unstimulated controls. Each paired dataset represents an individual experimental biological replicate. P values acquired by performing paired Student's t-test.

**S4F:** Quantification of data in **4A** depicting B cell nuclear p65, highlighting effect of high dose ionomcyin (1μM). gMFI values were normalized to unstimulated controls. Each paired dataset represents an individual experimental biological replicate. P values acquired by performing paired student's t-test.

**S4G:** Quantification of data in **S4I** depicting B cell nuclear NFAT2, highlighting effect of high dose ionomcyin (1μM). gMFI values were normalized to unstimulated controls. Each paired dataset represents an individual experimental biological replicate. P values acquired by performing paired student’s t-test.

**S4H:** pErk staining in MD4 B cells incubated with DGK inhibitor (R59949 100μM) for 30 minutes prior to stimulation with anti-IgM (10μg/mL) or PMA (20ng/mL). Data represents B220+ cells and is representative of two experiments.
**S4I:** p65 staining in isolated B and CD4+ T cell nuclei after incubation with DGK inhibitor (R59949 100μM) for 30 minutes and stimulation for 20 minutes.
**S4J:** NFAT2 staining in B cell nuclei. MD4 B cells were isolated from pooled splenocytes and lymph node cells and stimulated with PMA (20ng/mL), ionomycin (1μM), anti-IgM (10μg/mL), sHEL (1μg/mL) or pHEL (1pM) alone or with additional PMA (20, 2, 0.2 ng/mL) or ionomycin (1, 0.1, 0.01μM). Data representative of three experiments.
**S4K:** Quantification of data in **S4I**, highlighting effect of medium dose of PMA (20ng/mL). gMFI values were normalized to unstimulated controls. Each paired dataset represents an individual experimental biological replicate. P values acquired by performing paired Student’s t-test.
**S4L:** NFAT2 staining in isolated B and CD4+ T cell nuclei after incubation with DGK inhibitor (R59949 100μM) for 30 minutes and stimulation for 20 minutes.

**Supplementary Figure 5:**
**S5A:** CD69 staining of B cells were pre-incubated with R568 IRAK1/IRAK4 inhibitor (Rigel) prior to overnight stimulation. Data representative of two independent experiments.
**S5B:** Detailed PCA analysis between stimulation conditions using four distinct PCs. Labels highlight the top four loadings (transcripts) that drive divergence between PCs.
**S5C:** Transcript quantification of key drivers of PCA analysis from **S5B**.

**Supplementary Figure 6:**

**SGA:** Quantification of additional transcripts from RNAseq enriched in sHEL-stimulated B cells.
**S6B:** Quantification of *DGKa* and *DGKz* transcripts from RNAseq.
**S6C:** Bulk splenocytes and lymph nodes cells were treated with Cyclosporin A 1μM for 30 minutes prior to stimulation for 40hrs with respective stimuli. Histograms depict intracellular T-bet and surface PD-1 staining.
**S6D:** Quantification of gMFI data in **S6C**, normalized to unstimulated controls. P values acquired from Student's paired t-test.

**Supplementary Figure 7:**
**S7A:** GSEA performed to compare genes upregulated in sHEL+aCD40- (right) versus sHEL-stimulated B cells to the data set of hallmark Myc target genes (MM3887). NES = normalized enrichment score. FDRq = false discovery rate q-value.
**S7B-C:** GSEA performed to compare genes upregulated in pHEL- versus sHEL-stimulated B cells to the data set of Myc-dependent genes that are induced **(B)** or repressed **(C) b**y LPS stimulation (GSE126340). NES = normalized enrichment score. FDRq = false discovery rate q-value.
**S7D:** Isolated MD4 B cells were cultured for 72hrs with denoted stimuli. Graph depicts percentage of live B220+ cells. Each data point represents a biological replicate.
P values acquired from Student's paired t-test.
**S7E:** Isolated MD4 B cells were cultured for 72hrs with denoted stimuli with or without additional BAFF (20ng/mL). Graph depicts percentage of live B220+ cells. Each data series represents a biological replicate. P values acquired from Student's paired t-test.
**S7F:** Heatmap map showing expression of pro- and anti-apoptotic transcripts from RNAseq data.
**S7G:** Graphs depicting expression of notable pro- (top) and anti- (bottom) apoptotic transcripts from RNAseq data.
**S7H:** Isolated MD4 B cells were labeled with CTV and cultured for 72hrs with denoted stimuli with or without additional BAFF. Graph depicts proliferation index, calculated from CTV dilutions. Each data series represents a biological replicate.

**Supplementary Figure 8:**

**S8A:** HELD with C-terminal fusion of the Eα peptide (HELD-Eα) was cloned, expressed and purified to 97% purity as determined in a denaturing polycryladmide gel.

**S8B:** MD4 splenocytes were stimulated with Eα peptide (10, 1ug/mL), sHELD-Eα (1, 0.1, 0.01 μg/mL) or pHELD-Eα (10, 1, 0.1pM) overnight and stained for surface MHC II, CD86 and YAe expression. Stars denote matched doses of HEL protein between sHELD- Eα and pHELD- Eα.

**S8C:** Polyclonal CD4+ T cells were co-cultured with MD4 B cells and noted stimuli (PMA 20ng/mL + ionomycin 1uM, pHELT 10pM, pHELT-OVA 10pM, or OVA (5μM) overnight. Plots depict CD69 staining on CD4+ T cells denoted as live CD4+ Va2- Vb5- cells. Data representative of three experiments.

**S8D:** Model for divergent B cell responses to soluble and particulate antigen.

### Figure S1

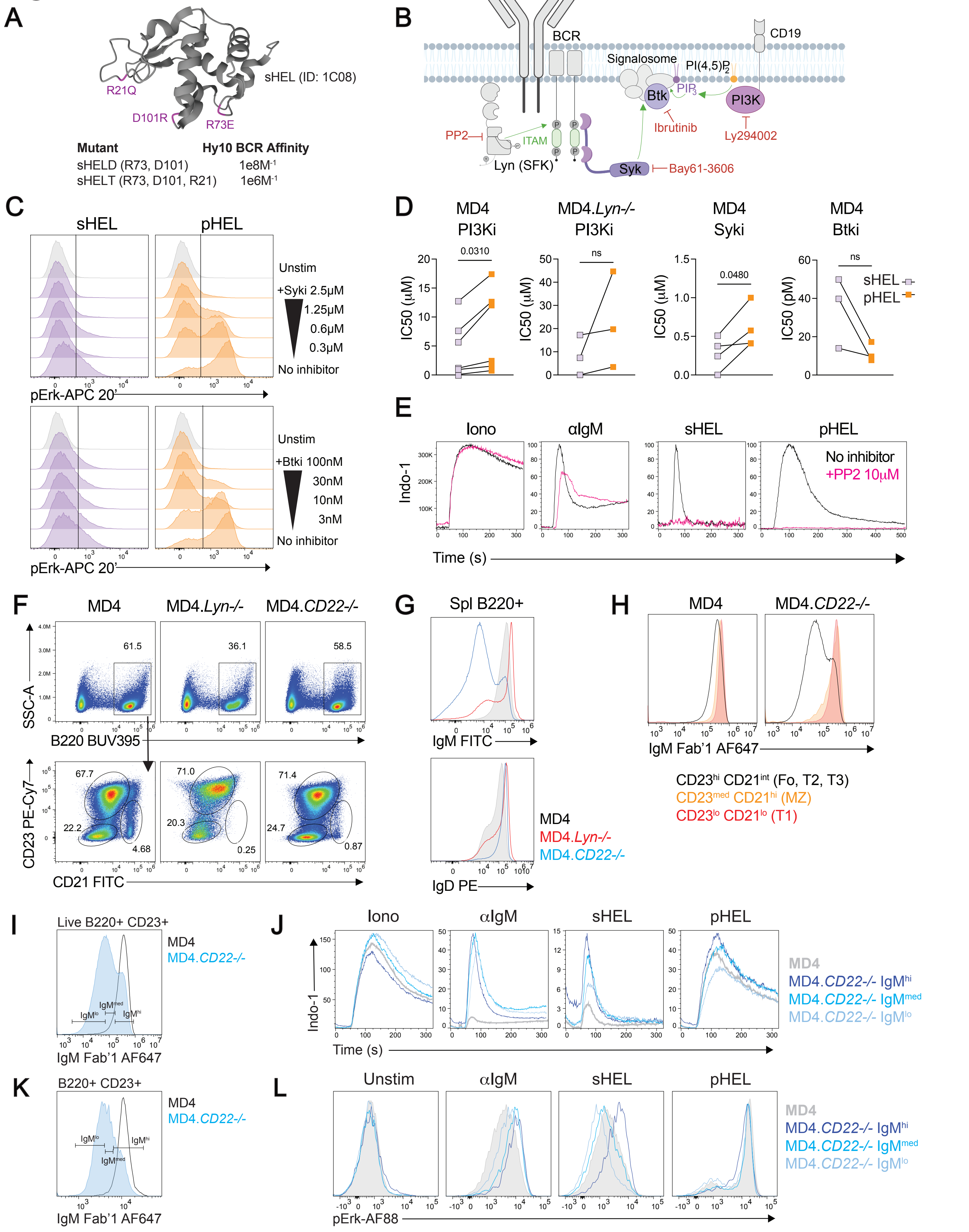

A
sHEL (ID: 1C08)
R21Q
D101R
R73E
Mutant
Hy10 BCR Affinity
sHELD (R73, D101) 1e8M-1
sHELT (R73, D101, R21) 1e6M-1
B
BCR
CD19
Signalosome
PI(4,5)P2
PIP3
Btk
PI3K
PP2
ITAM
Ibrutinib
Ly294002
Lyn (SFK)
Syk
Bay61-3606
C
sHEL
pHEL
Unstim
+Syki 2.5μM
1.25μM
0.6μM
0.3μM
No inhibitor
pErk-APC 20'
+Btki 100nM
30nM
10nM
3nM
D
MD4 PI3Ki
MD4.Lyn-/- PI3Ki
MD4 Syki
MD4 Btki
IC50 (μM)
IC50 (pM)
0.0310
ns
0.0480
sHEL
pHEL
E
Iono
αIgM
sHEL
pHEL
Indo-1
No inhibitor
+PP2 10μM
Time (s)
F
MD4
MD4.Lyn-/-
MD4.CD22-/-
SSC-A
B220 BUV395
61.5
36.1
58.5
CD23 PE-Cy7
CD21 FITC
67.7
22.2
4.68
71.0
20.3
0.25
71.4
24.7
0.87
G
Spl B220+
IgM FITC
IgD PE
MD4
MD4.Lyn-/-
MD4.CD22-/-
H
MD4
MD4.CD22-/-
IgM Fab'1 AF647
CD23hi CD21int (Fo, T2, T3)
CD23med CD21hi (MZ)
CD23lo CD21lo (T1)
I
Live B220+ CD23+
MD4
MD4.CD22-/-
IgMlo
IgMmed
IgMhi
IgM Fab'1 AF647
J
Iono
αIgM
sHEL
pHEL
Indo-1
Time (s)
MD4
MD4.CD22-/- IgMhi
MD4.CD22-/- IgMmed
MD4.CD22-/- IgMlo
K
B220+ CD23+
MD4
MD4.CD22-/-
IgMlo
IgMhi
IgMmed
IgM Fab'1 AF647
L
Unstim
αIgM
sHEL
pHEL
pErk-AF88
MD4
MD4.CD22-/- IgMhi
MD4.CD22-/- IgMmed
MD4.CD22-/- IgMlo

### Figure S2

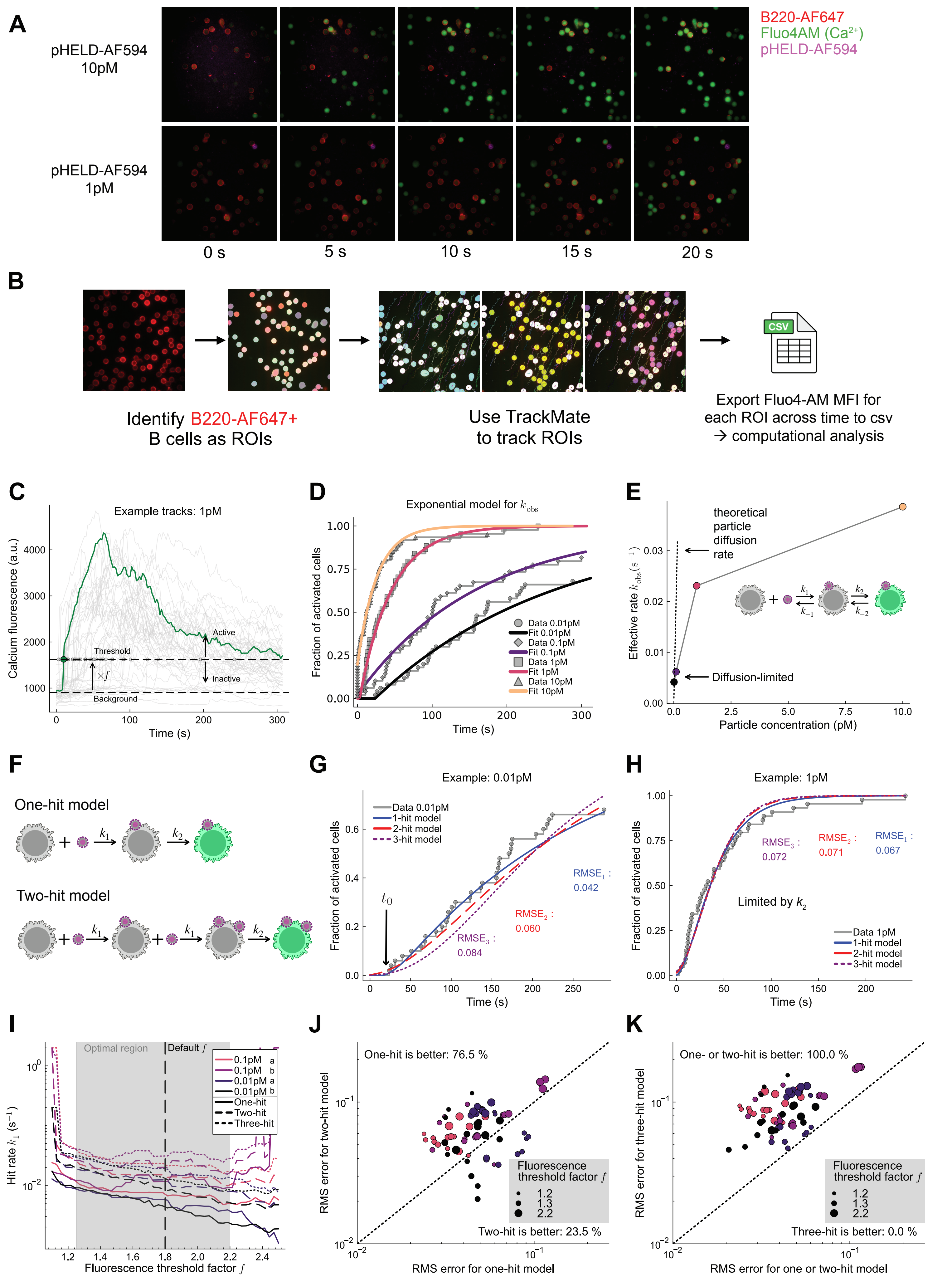
A
pHELD-AF594 10pM
pHELD-AF594 1pM
B220-AF647
Fluo4AM (Ca2+)
pHELD-AF594
0 s
5 s
10 s
15 s
20 s
B
Identify B220-AF647+ B cells as ROIs
Use TrackMate to track ROIs
CSV
Export Fluo4-AM MFI for each ROI across time to csv → computational analysis
C
Example tracks: 1pM
Calcium fluorescence (a.u.)
Threshold
Active
Inactive
Background
Time (s)
D
Exponential model for k_obs
Fraction of activated cells
Data 0.01pM
Fit 0.01pM
Data 0.1pM
Fit 0.1pM
Data 1pM
Fit 1pM
Data 10pM
Fit 10pM
Time (s)
E
theoretical particle diffusion rate
Effective rate k_obs (s−1)
Diffusion-limited
Particle concentration (pM)
F
One-hit model
Two-hit model
G
Example: 0.01pM
Data 0.01pM
1-hit model
2-hit model
3-hit model
RMSE1 : 0.042
RMSE2 : 0.060
RMSE3 : 0.084
Fraction of activated cells
Time (s)
H
Example: 1pM
RMSE3 : 0.072
RMSE2 : 0.071
RMSE1 : 0.067
Limited by k2
Data 1pM
1-hit model
2-hit model
3-hit model
Fraction of activated cells
Time (s)
I
Optimal region
Default f
0.1pM a
0.1pM b
0.01pM a
0.01pM b
One-hit
Two-hit
Three-hit
Hit rate k1 (s−1)
Fluorescence threshold factor f
J
One-hit is better: 76.5 %
Fluorescence threshold factor f
Two-hit is better: 23.5 %
RMS error for two-hit model
RMS error for one-hit model
K
One- or two-hit is better: 100.0 %
Fluorescence threshold factor f
Three-hit is better: 0.0 %
RMS error for three-hit model
RMS error for one or two-hit model

### Figure S3

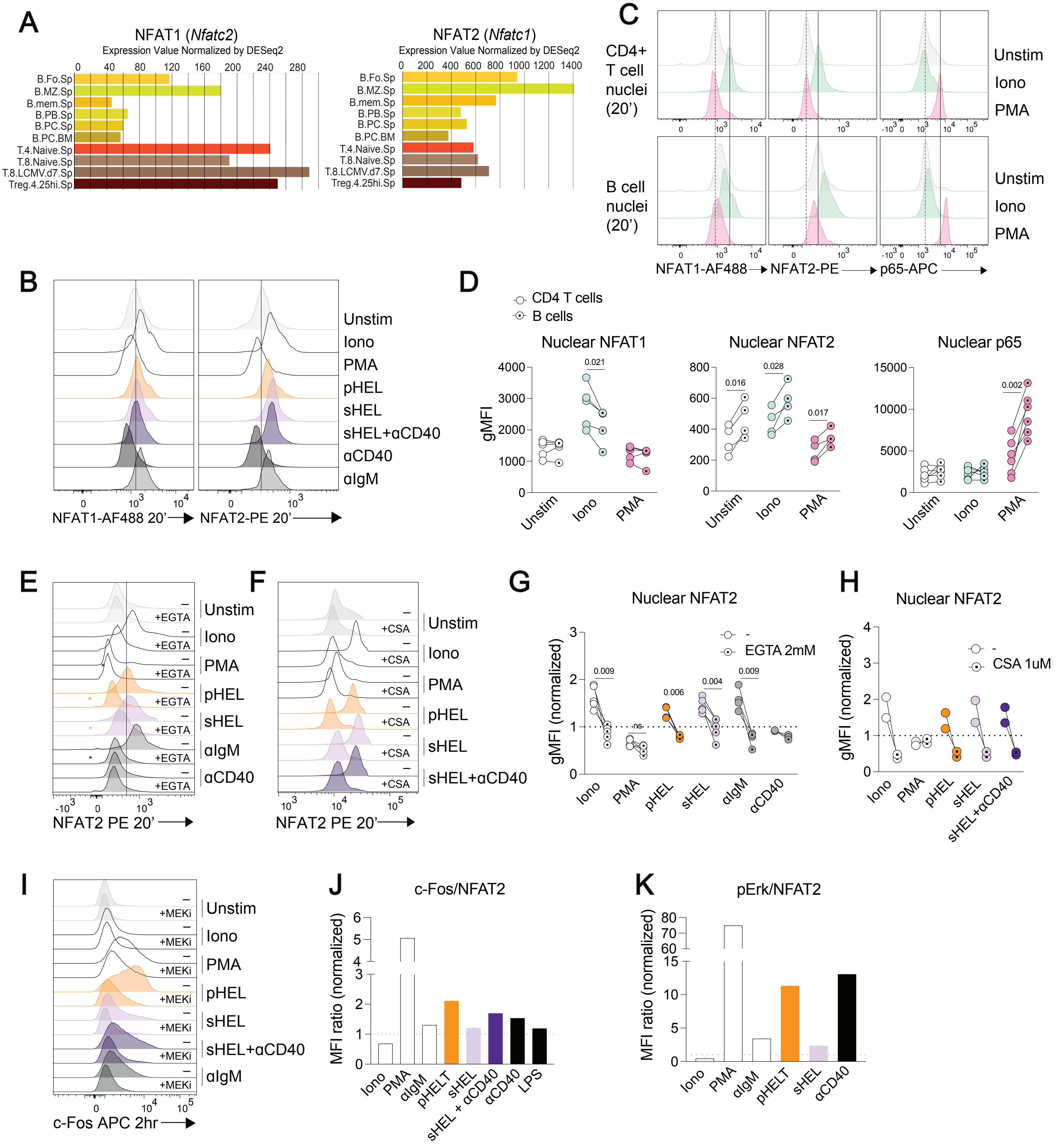

A
NFAT1 (Nfatc2)
Expression Value Normalized by DESeq2
NFAT2 (Nfatc1)
Expression Value Normalized by DESeq2
B.Fo.Sp
B.MZ.Sp
B.mem.Sp
B.PB.Sp
B.PC.Sp
B.PC.BM
T.4.Naive.Sp
T.8.Naive.Sp
T.8.LCMV.d7.Sp
Treg.4.25hi.Sp
C
CD4+ T cell nuclei (20')
B cell nuclei (20')
Unstim
Iono
PMA
NFAT1-AF488
NFAT2-PE
p65-APC
B
Unstim
Iono
PMA
pHEL
sHEL
sHEL+αCD40
αCD40
αIgM
NFAT1-AF488 20'
NFAT2-PE 20'
D
CD4 T cells
B cells
Nuclear NFAT1
Nuclear NFAT2
Nuclear p65
gMFI
E
+EGTA
NFAT2 PE 20'
F
+CSA
NFAT2 PE 20'
G
Nuclear NFAT2
EGTA 2mM
gMFI (normalized)
H
Nuclear NFAT2
CSA 1uM
gMFI (normalized)
I
+MEKi
c-Fos APC 2hr
J
c-Fos/NFAT2
MFI ratio (normalized)
Iono
PMA
αIgM
pHELT
sHEL
sHEL + αCD40
αCD40
LPS
K
pErk/NFAT2
MFI ratio (normalized)

### Figure S4

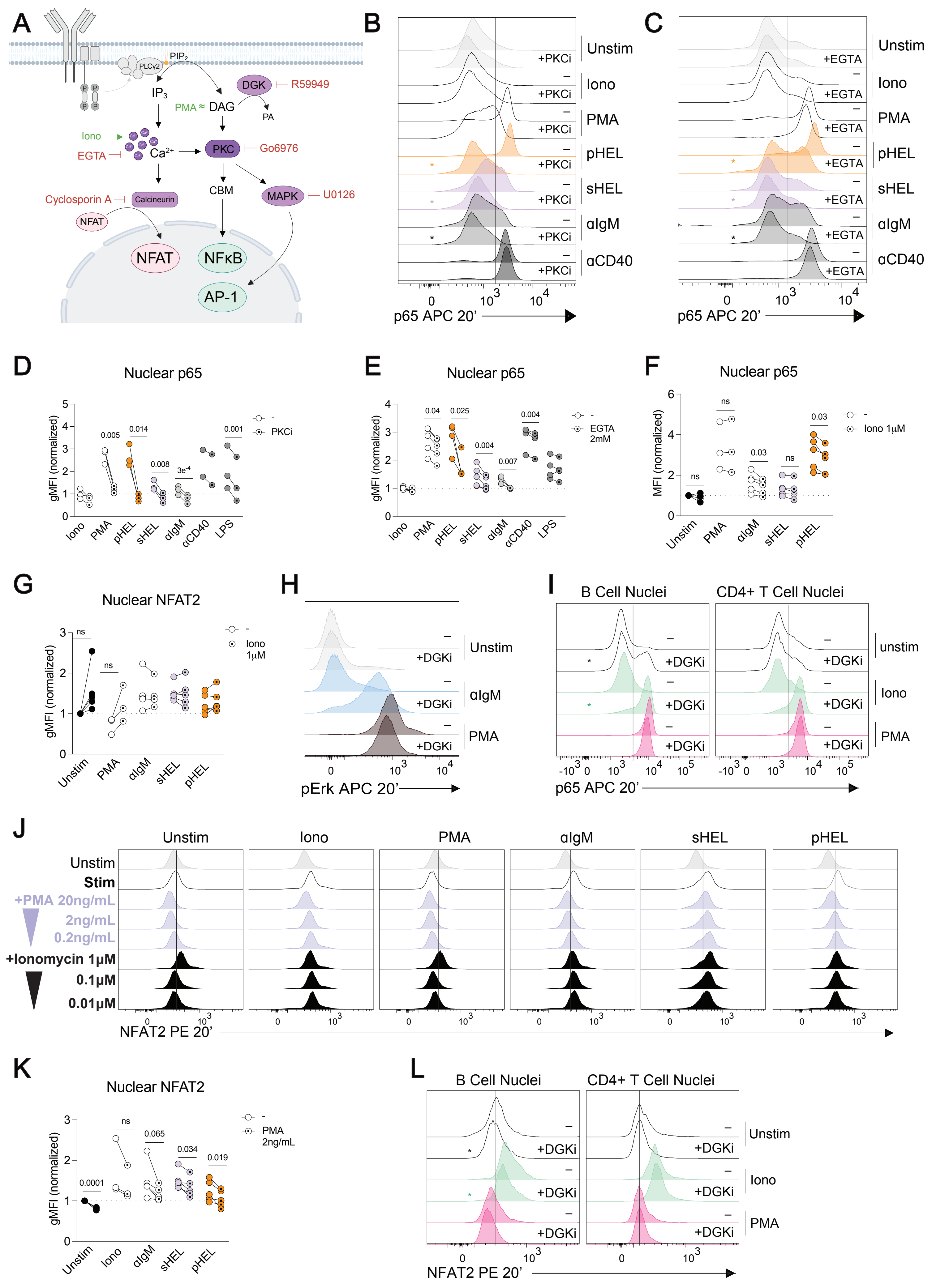

A
PIP2
PLCγ2
DGK
R59949
IP3
PMA ≈ DAG
PA
Iono
EGTA
Ca2+
PKC
Go6976
CBM
Cyclosporin A
Calcineurin
MAPK
U0126
NFAT
NFAT
NFκB
AP-1
B
Unstim
+PKCi
Iono
PMA
pHEL
sHEL
αIgM
αCD40
p65 APC 20'
C
Unstim
+EGTA
Iono
PMA
pHEL
sHEL
αIgM
αCD40
p65 APC 20'
D
Nuclear p65
gMFI (normalized)
0.005
0.014
0.008
3e-4
0.001
- PKCi
Iono PMA pHEL sHEL αIgM αCD40 LPS
E
Nuclear p65
gMFI (normalized)
0.04
0.025
0.004
0.007
0.004
- EGTA 2mM
Iono PMA pHEL sHEL αIgM αCD40 LPS
F
Nuclear p65
MFI (normalized)
ns
ns
0.03
ns
0.03
- Iono 1μM
Unstim PMA αIgM sHEL pHEL
G
Nuclear NFAT2
gMFI (normalized)
ns
ns
- Iono 1μM
Unstim PMA αIgM sHEL pHEL
H
Unstim
+DGKi
αIgM
PMA
pErk APC 20'
I
B Cell Nuclei
CD4+ T Cell Nuclei
unstim
+DGKi
Iono
PMA
p65 APC 20'
J
Unstim
Iono
PMA
αIgM
sHEL
pHEL
Unstim
Stim
+PMA 20ng/mL
2ng/mL
0.2ng/mL
+Ionomycin 1μM
0.1μM
0.01μM
NFAT2 PE 20'
K
Nuclear NFAT2
gMFI (normalized)
0.0001
ns
0.065
0.034
0.019
- PMA 2ng/mL
Unstim Iono αIgM sHEL pHEL
L
B Cell Nuclei
CD4+ T Cell Nuclei
Unstim
+DGKi
Iono
PMA
NFAT2 PE 20'

### Figure S5

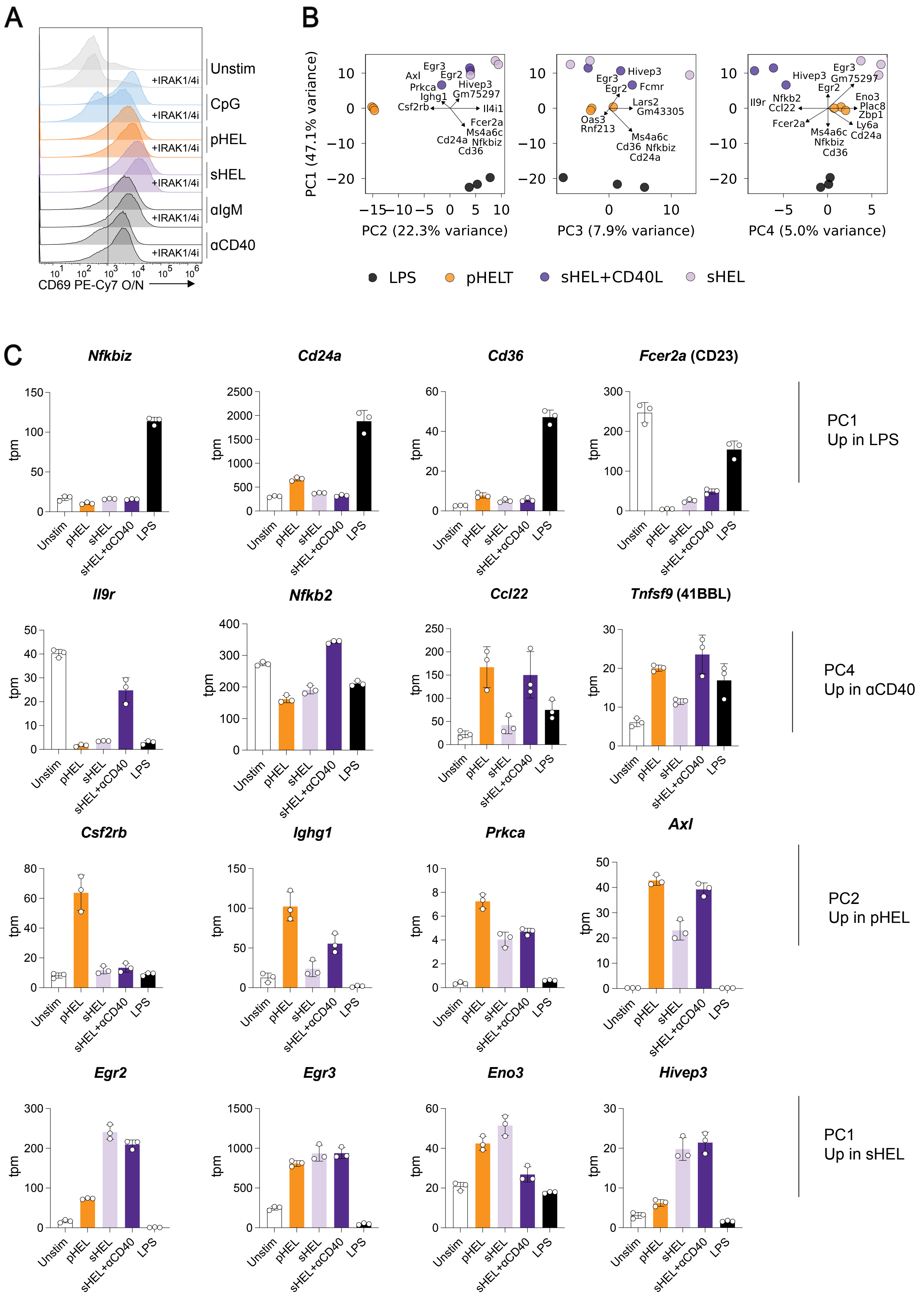

A
Unstim
CpG
pHEL
sHEL
αIgM
αCD40
+IRAK1/4i
CD69 PE-Cy7 O/N
B
PC1 (47.1% variance)
PC2 (22.3% variance)
PC3 (7.9% variance)
PC4 (5.0% variance)
Egr3
Axl
Egr2
Prkca
Hivep3
Ighg1
Gm75297
Csf2rb
Il4i1
Fcer2a
Ms4a6c
Cd24a
Nfkbiz
Cd36
Fcmr
Lars2
Gm43305
Oas3
Rnf213
Il9r
Nfkb2
Ccl22
Eno3
Plac8
Zbp1
Ly6a
LPS
pHELT
sHEL+CD40L
sHEL
C
Nfkbiz
Cd24a
Cd36
Fcer2a (CD23)
PC1
Up in LPS
Il9r
Nfkb2
Ccl22
Tnfsf9 (41BBL)
PC4
Up in αCD40
Csf2rb
Ighg1
Prkca
Axl
PC2
Up in pHEL
Egr2
Egr3
Eno3
Hivep3
PC1
Up in sHEL
tpm
Unstim
pHEL
sHEL
sHEL+αCD40
LPS

### Figure S6

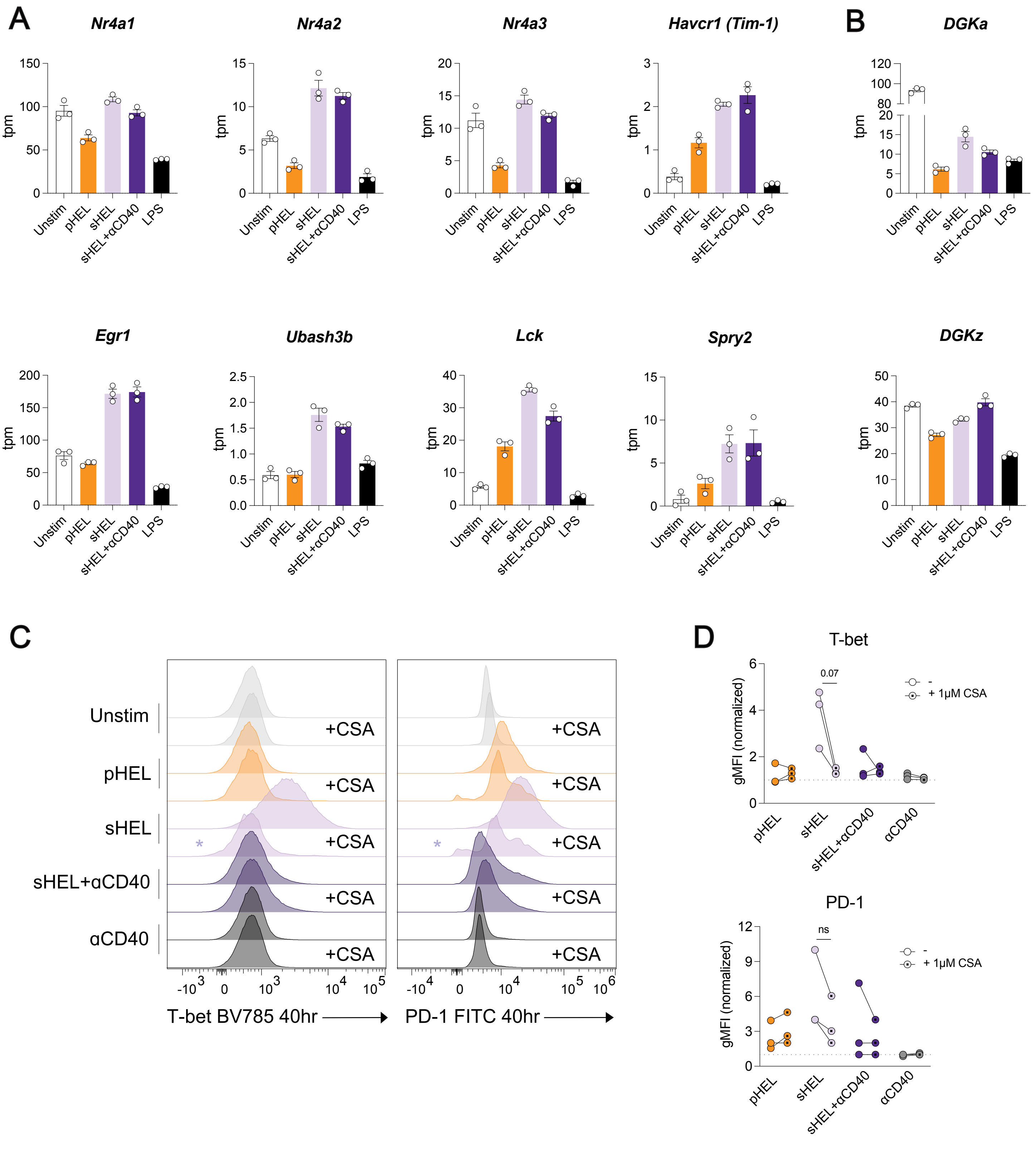
A
Nr4a1
Nr4a2
Nr4a3
Havcr1 (Tim-1)
B
DGKa
Egr1
Ubash3b
Lck
Spry2
DGKz
tpm
Unstim
pHEL
sHEL
sHEL+αCD40
LPS
C
D
T-bet
PD-1
+CSA
*
T-bet BV785 40hr
PD-1 FITC 40hr
gMFI (normalized)
0.07
ns
-
+ 1μM CSA
αCD40

### Figure S7

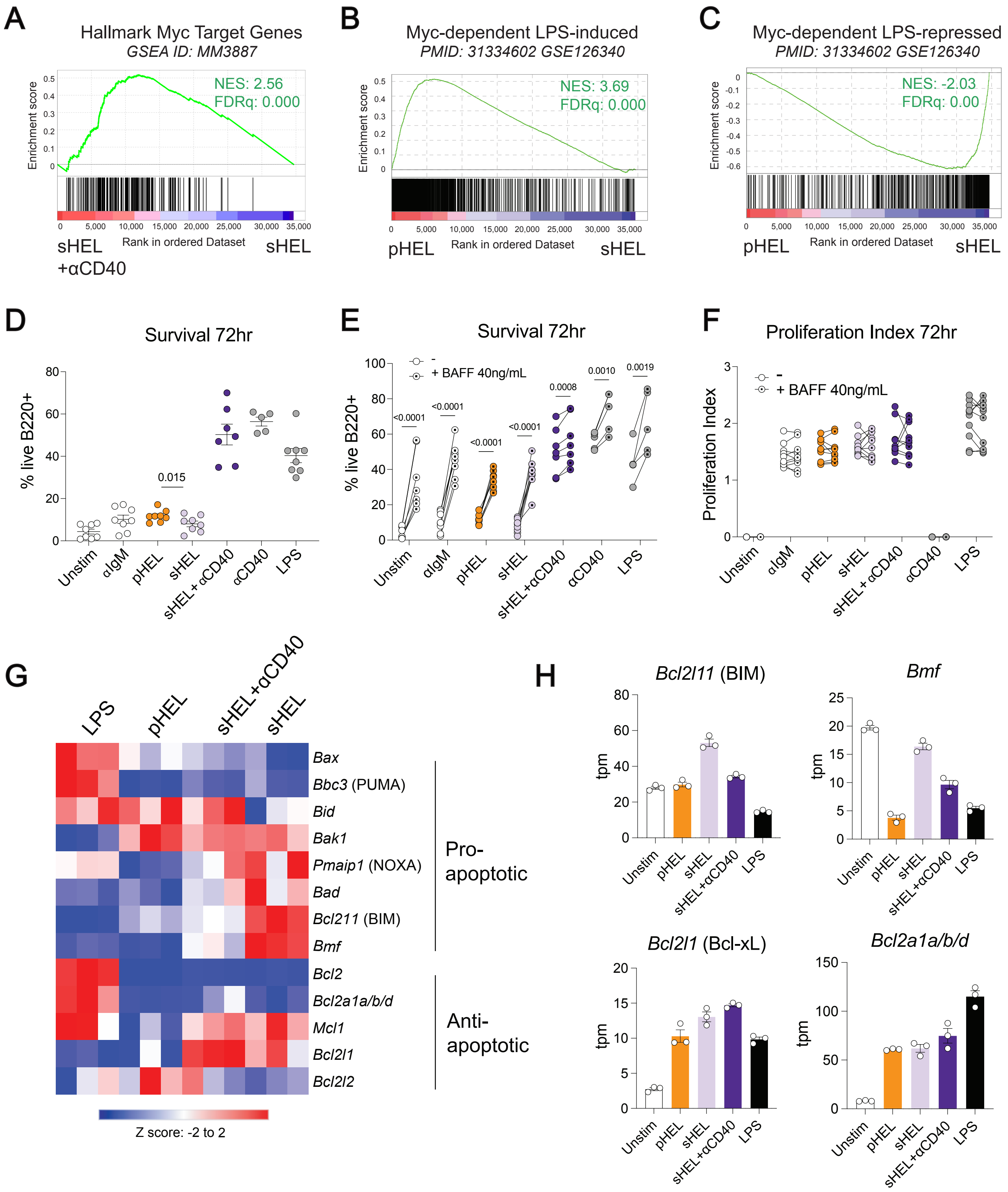

A
Hallmark Myc Target Genes
GSEA ID: MM3887
NES: 2.56
FDRq: 0.000
Enrichment score
Rank in ordered Dataset
sHEL +αCD40
sHEL
B
Myc-dependent LPS-induced
PMID: 31334602 GSE126340
NES: 3.69
FDRq: 0.000
pHEL
sHEL
C
Myc-dependent LPS-repressed
PMID: 31334602 GSE126340
NES: -2.03
FDRq: 0.00
pHEL
sHEL
D
Survival 72hr
% live B220+
0.015
Unstim
αIgM
pHEL
sHEL
sHEL + αCD40
αCD40
LPS
E
Survival 72hr
-
+ BAFF 40ng/mL
<0.0001
<0.0001
<0.0001
<0.0001
0.0008
0.0010
0.0019
F
Proliferation Index 72hr
Proliferation Index
G
LPS
pHEL
sHEL+αCD40
sHEL
Bax
Bbc3 (PUMA)
Bid
Bak1
Pmaip1 (NOXA)
Bad
Bcl211 (BIM)
Bmf
Bcl2
Bcl2a1a/b/d
Mcl1
Bcl2l1
Bcl2l2
Pro-apoptotic
Anti-apoptotic
Z score: -2 to 2
H
Bcl2l11 (BIM)
Bmf
Bcl2l1 (Bcl-xL)
Bcl2a1a/b/d
tpm
Unstim
pHEL
sHEL
sHEL+αCD40
LPS

### Figure S8

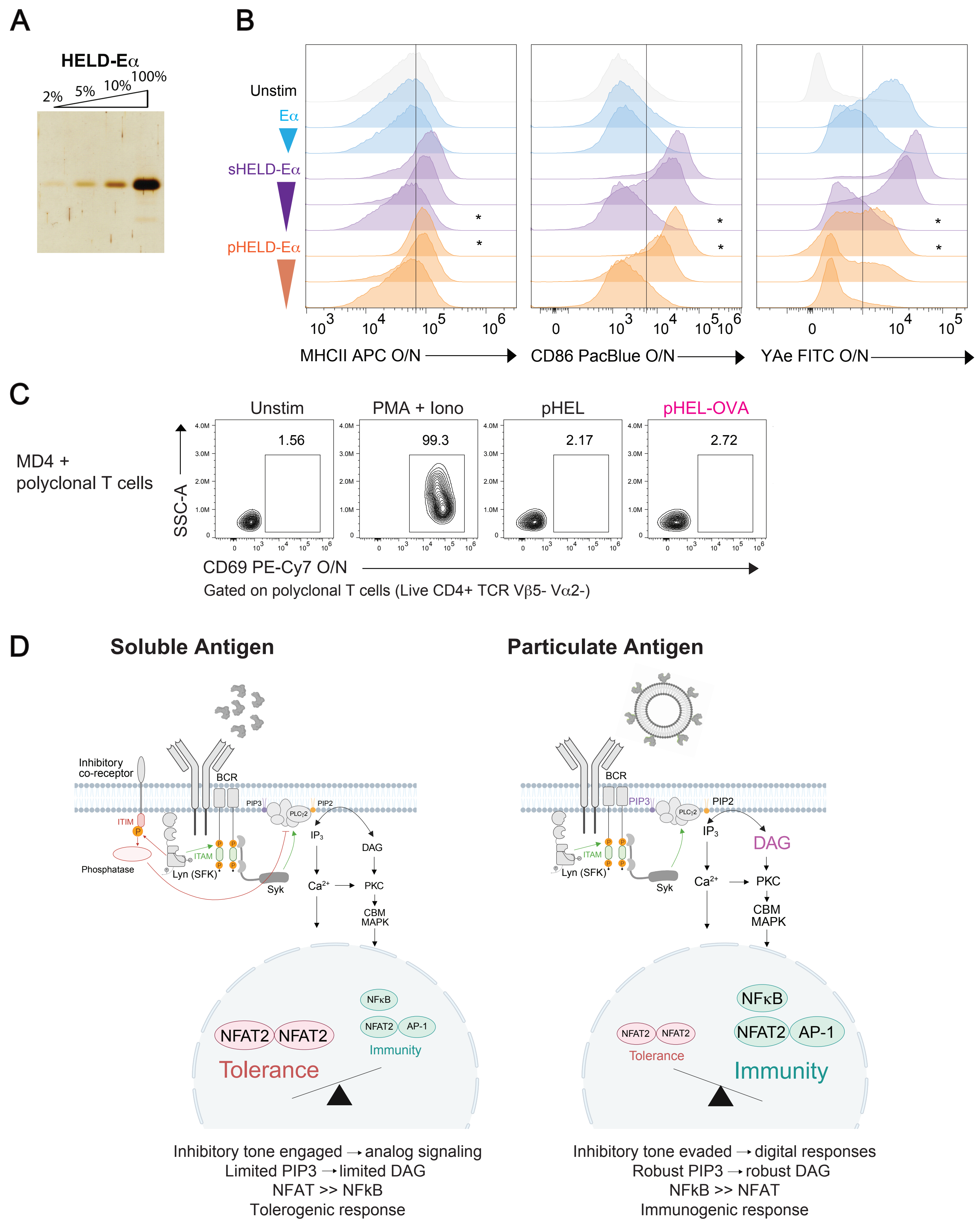
A
HELD-Eα
2%
5%
10%
100%
B
Unstim
Eα
sHELD-Eα
pHELD-Eα
MHCII APC O/N
CD86 PacBlue O/N
YAe FITC O/N
C
MD4 +
polyclonal T cells
Unstim
PMA + Iono
pHEL
pHEL-OVA
1.56
99.3
2.17
2.72
SSC-A
CD69 PE-Cy7 O/N
Gated on polyclonal T cells (Live CD4+ TCR Vβ5- Vα2-)
D
Soluble Antigen
Particulate Antigen
Inhibitory
co-receptor
BCR
ITIM
Phosphatase
ITAM
Lyn (SFK)
Syk
PIP3
PIP2
PLCγ2
IP3
DAG
Ca2+
PKC
CBM
MAPK
NFκB
NFAT2
AP-1
Immunity
Tolerance
Inhibitory tone engaged → analog signaling
Limited PIP3 → limited DAG
NFAT >> NFkB
Tolerogenic response
Inhibitory tone evaded → digital responses
Robust PIP3 → robust DAG
NFkB >> NFAT
Immunogenic response
